# Discovery of the fastest myosin, its amino acid sequence, and structural features

**DOI:** 10.1101/2021.05.06.442907

**Authors:** Takeshi Haraguchi, Masanori Tamanaha, Kano Suzuki, Kohei Yoshimura, Takuma Imi, Motoki Tominaga, Hidetoshi Sakayama, Tomoaki Nishiyama, Takeshi Murata, Kohji Ito

**Affiliations:** Department of Biology, Graduate School of Science, Chiba University, Chiba 263-8522, Japan; Department of Chemistry, Graduate School of Science, Chiba University, Chiba 263-8522, Japan; Faculty of Education and Integrated Arts and Sciences, Waseda University, 2-2 Wakamatsu-cho, Shinjuku-ku, Tokyo 162-8480, Japan; Major in Integrative Bioscience and Biomedical Engineering, Graduate School of Science and Engineering, Waseda University, 2-2 Wakamatsu-cho, Shinjuku-ku, Tokyo 162-8480, Japan; Department of Biology, Graduate School of Science, Kobe University, 1-1 Rokkodai, Nada-ku, Kobe 657-8501, Japan; Research Center for Experimental Modeling of Human Disease, Kanazawa University, Kanazawa 920-0934, Japan; Molecular Chirality Research Center, Chiba University, Chiba 263-8522, Japan

**Keywords:** myosin XI, X-ray crystallography, cytoplasmic streaming, actin

## Abstract

Cytoplasmic streaming with extremely high velocity (~70 μm s^−1^) occurs in cells of the characean algae (*Chara*). Because cytoplasmic streaming is caused by organelle-associated myosin XI sliding along actin filaments, it has been suggested that a myosin XI, which has a velocity of 70 μm s^−1^, the fastest myosin measured so far, exists in *Chara* cells. However, the previously cloned *Chara corallina* myosin XI (*Cc*XI) moved actin filaments at a velocity of around 20 μm s^−1^, suggesting that an unknown myosin XI with a velocity of 70 μm s^−1^ may be present in *Chara*. Recently, the genome sequence of *Chara braunii* has been published, revealing that this alga has four myosin XI genes. In the work reported in this paper, we cloned these four myosin XIs (*Cb*XI-1, 2, 3, and 4) and measured their velocities. While the velocities of *Cb*XI-3 and *Cb*XI-4 were similar to that of *Cc*XI, the velocities of *Cb*XI-1 and *Cb*XI-2 were estimated to be 73 and 66 μm s^−1^, respectively, suggesting that *Cb*XI-1 and *Cb*XI-2 are the main contributors to cytoplasmic streaming in *Chara* cells and showing that *Cb*XI-1 is the fastest myosin yet found. We also report the first atomic structure (2.8 Å resolution) of myosin XI using X-ray crystallography. Based on this crystal structure and the recently published cryo-EM structure of acto-myosin XI at low resolution (4.3 Å), it appears that the actin-binding region contributes to the fast movement of *Chara* myosin XI. Mutation experiments of actin-binding surface loop 2 support this hypothesis.

**Significance statement:** It has been suggested for more than 50 years that the fastest myosin in the biological world, with a velocity of 70 μm s^−1^, exists in the alga *Chara* because cytoplasmic streaming with a velocity of 70 μm s^−1^ occurs in *Chara* cells. However, a myosin with that velocity has not yet been identified. In this work, we succeeded in cloning a myosin XI with a velocity of 73 μm s^−1^, the fastest myosin so far measured. We also successfully crystallized myosin XI for the first time. Structural analyses and mutation experiments suggest that the central regions that define the fast movement of *Chara* myosin XI are the actin-binding sites.

## Introduction

Myosins are motor proteins that convert chemical energy, ATP, to physical force to move actin filaments. Phylogenetic analyses of myosin motor domain (MD) sequences have shown that there are at least 79 myosin classes, with several subclasses under each class (1). Myosins of different classes and subclasses differ significantly in properties such as velocity, ATPase activity, and duty ratio (the proportion of the ATPase cycle in which the MD remains strongly bound to actin) and perform different intracellular functions (2). The diversity of properties of these classes and subclasses arise from differences in the rates of the binding and dissociation of ATP, ADP, and actin filaments (3).

Plants have two plant-specific myosin classes, myosin VIII and myosin XI. Myosin VIII moves actin filaments at very slow velocities (4) and is involved in endocytosis, cell plate formation, and plasmodesmatal functioning in plants (5–7). Myosin XI produces an intracellular flow known as cytoplasmic streaming in plant cells by moving on actin filaments while binding organelles via its tail domain. Cytoplasmic streaming facilitates the distribution of molecules and vesicles throughout large plant cells (8–12). The velocities of myosin XI are generally high, and the molecule specializes in cytoplasmic streaming. Some cells of characean algae (*Chara*) are very large, being up to 10 cm long and 0.1 cm in diameter. Very fast cytoplasmic streaming, of up to 70 μm s^−1^, is required for the dispersal of molecules and vesicles into the giant *Chara* cells (13).

Based on the velocity of cytoplasmic streaming in *Chara* cells, it has long been suggested that *Chara* has a myosin moving on actin filaments at 70 μm s^−1^ (13–17). This velocity is 10 times faster than the velocity of fast skeletal muscle myosin and the fastest of all myosins measured. The development of approaches for cloning this ultrafast myosin are urgently needed. Details of the sequence of the protein and the ability to work with cloned myosin constructs will allow the investigation of the mechanisms that control the myosin velocity and facilitate investigation of the detailed chemical-mechanical conversion mechanism of myosin (18). Kashiyama et al. cloned the cDNA of *Chara* myosin from a *Chara corallina* cDNA library by immunoscreening, using antibodies against purified *Chara corallina* myosin (19). Morikawa et al. also cloned the cDNA of *Chara* myosin using the same method as that used by Kashiyama et al. (20). The sequences of the MD of myosins cloned by the two groups were identical, and there was a 15 amino acid indel variation in the tail domain, a finding that indicates potential alternative splicing in the tail domain. The *Chara corallina* myosin XI (*Cc*XI) contains six IQ motifs, which are light chain binding sites. It was not possible to express the protein and measure its velocity using the cloned *Cc*XI, because the myosin light chains that bind to the six IQ motifs of *Cc*XI have not been identified. Therefore, the functional expression of *Cc*XI has been carried out using either a *Cc*XI MD construct that did not have the myosin light chain binding sites (IQ motifs) or chimeric full-length *Cc*XI constructs in which IQ motifs and myosin light chains of *Cc*XI were replaced with those of other myosins. The velocity of *Cc*XI was then estimated from the velocity measured using these constructs. The estimated velocity of *Cc*XI was about 20 μm s^−1^ or less at 25 °C (10, 21–25), which is less than about one-third of the velocity of cytoplasmic streaming observed in *Chara* cells. Three possibilities have been suggested as to why the velocities of *Cc*XI obtained using the recombinant constructs were different from that expected from cytoplasmic streaming. (1) The recombinant *Cc*XI constructs do not have the same IQ motifs and myosin light chains as native *Cc*XI, and this substitution may have affected the velocity. (2) *Cc*XI may undergo a post-translational modification in *Chara* cells, which may increase the velocity of *Cc*XI in cells. (3) A myosin XI gene other than *Cc*XI may be present in *Chara* cells, and this myosin XI may be responsible for cytoplasmic streaming with a velocity of 70 μm s^−1^.

Recently, a genome project has been conducted for *Chara braunii* (26). *Chara braunii* is phylogenetically close to *Chara corallina* (27–29), and both species have the same cytoplasmic streaming velocity, 70 μm s^−1^. The *Chara* genome project revealed that the *Chara braunii* genome contains four myosin XI genes.

In this study, we cloned the four *Chara braunii* myosin XIs and named them *Cb*XI-1, *Cb*XI-2, *Cb*XI-3, and *Cb*XI-4. Phylogenetic analyses indicated that the myosin XIs in *Chara* form a clade in streptophyte myosin XIs, expanded independently from seed plant myosin XIs, and gave rise to the four members in *Chara braunii. Cb*XI-4 may be an ortholog of *Cc*XI. We showed that the velocities of *Cb*XI-1 and *Cb*XI-2 were almost the same as the velocity of cytoplasmic streaming in *Chara* cells (70 μm s^−1^) and that *Cb*XI-1 is the fastest myosin in the biological world (73 μm s^−1^) and loop 2 in *Cb*XI-1 plays a vital role in the velocity. We also succeeded in crystallizing *Arabidopsis* myosin XI-2 (*At*XI-2), the first atomic structure of myosin XI, and a valuable comparator for the *Chara* myosin. Structural analyses and mutation experiments suggest that the central regions that define *Chara* myosin XI’s fast movement are the actin-binding sites.

## Results

### Phylogenetic relationships of the four *Chara braunii* myosin XIs

Until 2018, the only known myosin sequence in the genus *Chara* was that of *Cc*XI, which was cloned by Kashiyama (19) and Morimatsu (20) independently in 2000. Their results suggested that *Chara* has only one myosin XI gene. However, the *Chara braunii* genome project, published in 2018, revealed four myosin XI genes having intact motor domain: g50407, g48390, g24025, and g48658 (26). We named g50407, g48390, g24025, and g48658 as *Cb*XI-1, *Cb*XI-2, *Cb*XI-3, and *Cb*XI-4, respectively. The originally annotated g48658 (*Cb*XI-4) was truncated at the N-terminal 743 amino acids. The rest of the mRNA sequence of *Cb*XI-4 was identified on another scaffold, based on transcriptome assemblies (*SI Appendix*, *Materials and Methods* and accession nos: BR001749 and BR001750 for two isoforms). The full-length amino acid sequences of *Cb*XI-1, *Cb*XI-2, *Cb*XI-3, and *Cb*XI-4 are shown in *SI Appendix, Supplementary Text*. A schematic diagram of the *Cb*XIs deduced from the amino acid sequences is shown in Fig. 1. *Cb*XIs have typical domain structures of myosin XI.

**Figure 1.**
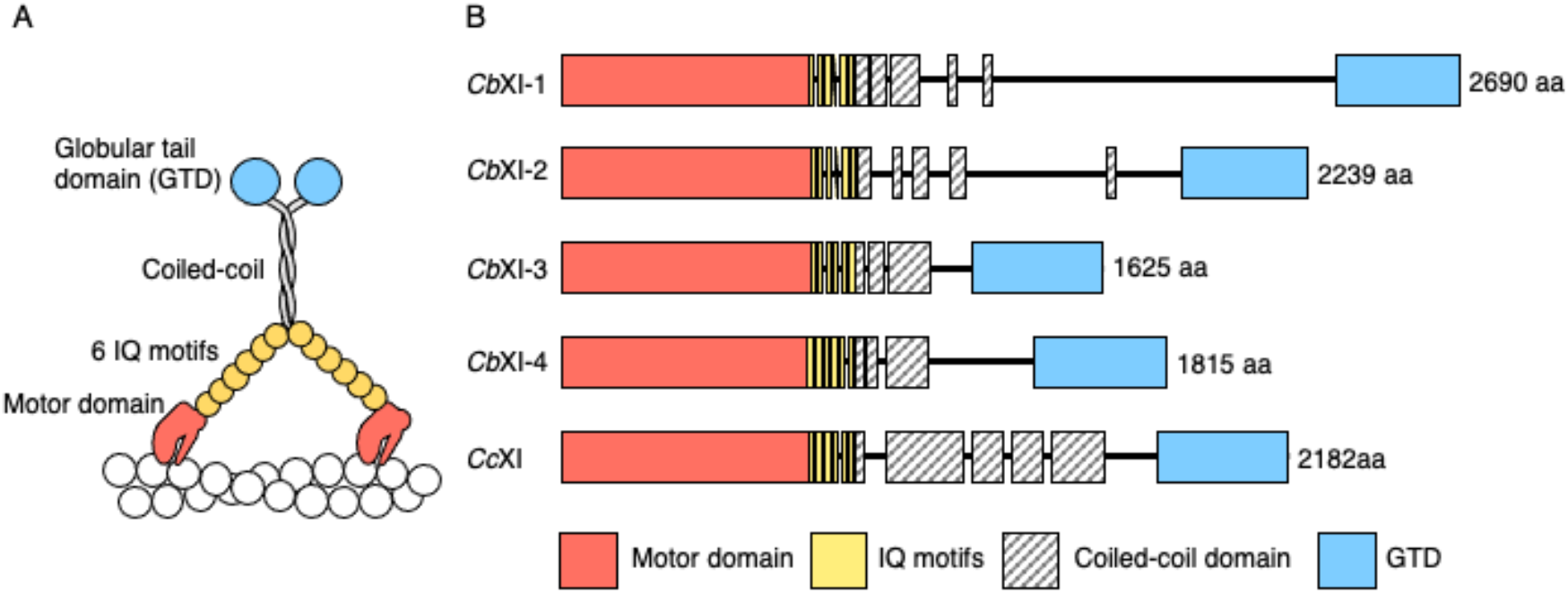
*Chara braunii* myosin XI structure deduced from its amino acid sequence. (*A*) Schematic diagrams of native *Chara braunii* myosin XIs (*CbXIs*). *CbXIs* show typical domain structures of myosin XI. They contain a motor domain (MD) with nucleotide and actin-binding sites, six IQ motifs to which six myosin light chains bind, an α-helical coiled-coil domain leading to dimer formation, and a globular tail domain (GTD). (*B*) Domain structures of four *Chara braunii* and one *Chara corallina* myosin XIs. Domains and motifs indicated by colored boxes were predicted using the MOTIF Search (https://www.genome.jp/tools/motif/) and COILS programs (43). Black lines are regions that were not recognized as known domains or motif structures. The length of each box and each line is shown to be proportional to the number of amino acids occupying each region.

We examined the phylogenetic relationships among the myosin XIs from *Chara* and representative green plants (Fig. 2). The phylogenetic tree indicated that streptophyte myosin XIs formed a well-supported clade including genes from *Klebsormidium nitens* and the Phragmoplastophyta, which includes *Chara*, *Spirogloea*, and the land plants. Though, the basal relationship within Phragmoplastophyta was not clearly resolved. The four *Chara braunii* myosin XI genes and a *Chara corallina* myosin XI gene formed a well-supported clade (Fig. 2. light-yellow box, Charales Myosin XI). Within the Charales myosin XI clade, *Cb*XI-1 and *Cb*XI-2 formed a clade (subgroup 1) sister to the remaining three genes (subgroup 2). *Cb*XI-4 and *Cc*XI formed a clade, and *Cb*XI-3 diverged earlier. The subgroup 1 have notably longer branches compared with subgroup 2 or other green plant myosin XIs. The proteins encoded by these two subgroups 1 genes are apparently larger than the subgroup 2 genes, especially long are the regions between coiled-coil and GTD domains (Fig. 1 *B*).

**Figure 2.**
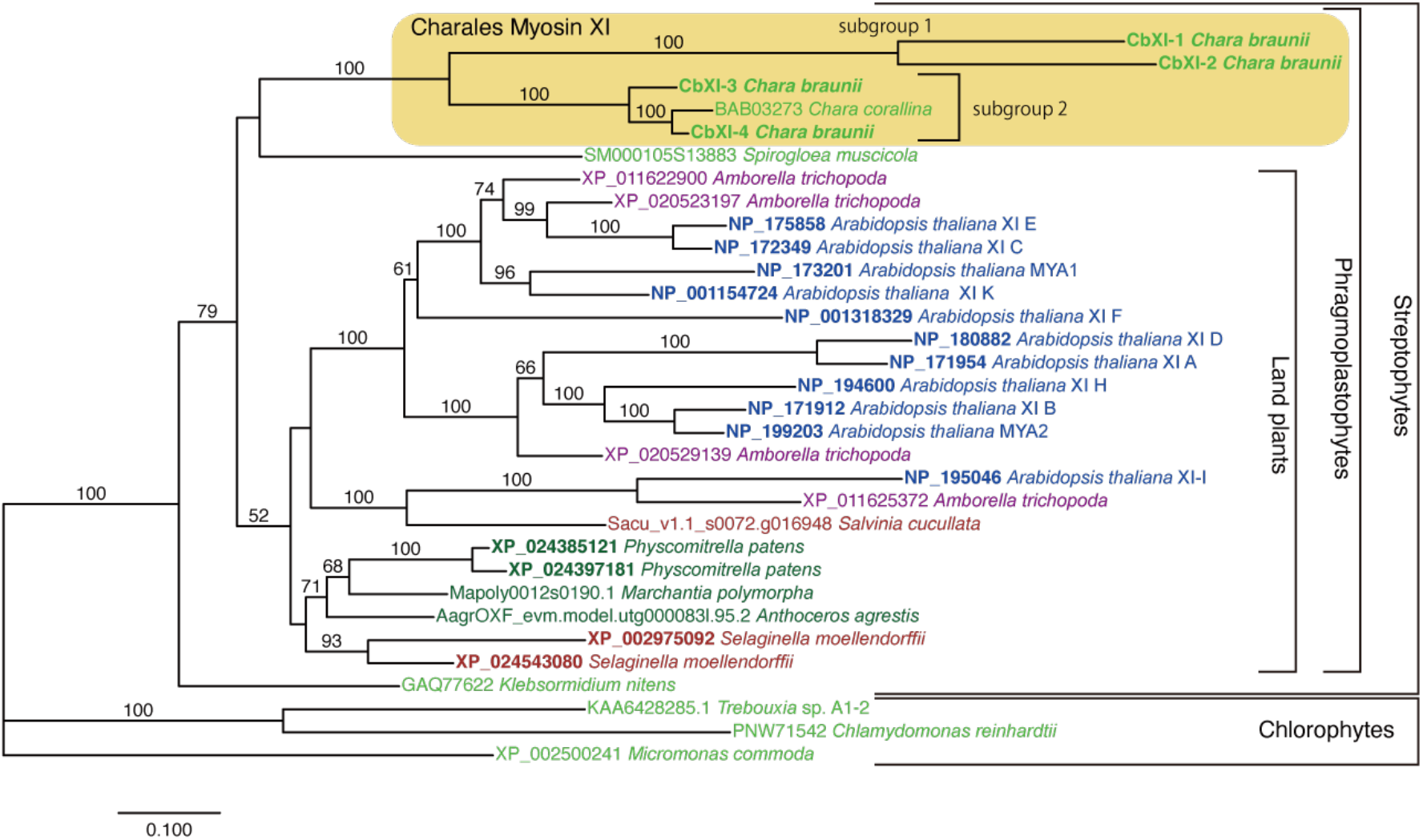
Phylogenetic relationship of green plant myosin XI genes. The phylogenetic tree was constructed under the L+GF model using RAxML. Amino acid sequences of 32 representative myosin XI genes including 5 genes from *Chara* were retained in the alignment and 1,060 sites in conserved regions were used for the analysis. Bootstrap analysis was performed with 1,000 replicate and the percentage values are indicated on each branch > 50%. The horizontal branch lengths are proportional to the estimated number of substitutions per site. Identifiers for *Salvinia cucullata*, *Anthoceros agrestis*, and *Marchantia polymorpha* are from respective genome databases; other identifiers are accession numbers for INSDC.

### Recombinant constructs of *Chara braunii* myosin XIs

To clarify the biochemical properties of the four *Cb*XIs, it was necessary to isolate and purify each *Cb*XI. However, it is very difficult to purify active myosins from *Chara* cells because most of the cell volume of *Chara* cells is occupied by the vacuole, which is rich in proteolytic enzymes, and the cytoplasm is only a small volume (30). Furthermore, it is virtually impossible to purify each of the four myosin XIs with similar molecular properties from *Chara braunii* cells. The only way to obtain each of four *Cb*XIs is to express and purify them using a recombinant construct. Using baculovirus expression systems for the functional expression of myosins with IQ motifs, co-expression with myosin light chains that bind to the IQ motifs is required. However, the myosin light chains of *Chara* myosin XIs have not been identified. Calmodulin binds IQ motifs and functions as myosin light chains for many unconventional myosins, such as myosin I, myosin V, and myosin VI. However, calmodulin did not function as myosin light chains for *Chara* myosins.

We therefore expressed *Cb*XIs using approaches that did not include the sequences of the IQ motifs of *Chara* myosins (See Fig.1*A* for the IQ motifs). We used two types of constructs: a *Cb*XI MD construct without IQ motifs (*Cb*XI MD) and a chimeric full-length *Cb*XI construct including the MD of *Cb*XI and six IQ motifs, coiled-coil, and tail domains of *Arabidopsis* myosin XI-2 (Chimeric *Cb*XI). Chimeric *Cb*XI constructs were co-expressed with *Arabidopsis* calmodulin, which is known to bind to the IQ motifs of *Arabidopsis* myosin XI-2 and function as myosin light chains (10). These constructs were expressed in a baculovirus system and purified by nickel-affinity and FLAG-affinity resins (*SI Appendix*, *Materials and Methods*).

### Velocities of *Chara braunii* myosin XIs

The sliding velocities of actin filaments by *Cb*XIs MD were measured using an antibody-based version of the *in vitro* motility assay at 25 °C. *Cb*XI-3 MD and *Cb*XI-4 MD moved actin filaments at velocities of 3.0 ± 0.2 and 3.1 ± 0.3 μm s^−1^, respectively, similar to that of *Cc*XI MD (22, 23). The velocities of *Cb*XI-1 MD and *Cb*XI-2 MD were 15 ± 0.7 μm s^−1^ and 13 ± 0.6 μm s^−1^, respectively, which were about threefold faster than that of *Cc*XI MD (22, 23) (Table 1). Actin velocities generated by myosins are approximately proportional to the lever arm length of myosin if the motor region is the same (31, 32). We have previously shown that this relationship between lever arm length and actin velocities by myosin generally holds for myosin XIs. The lever arm length of myosin XI MD is 1/6.6 that of full-length myosin XI (native myosin XI) containing six IQ motifs, and the velocity of myosin XI MD was 1/5 that of full-length myosin XI (33). Therefore, multiplying the velocity of myosin XI MD by a factor of 5 gives the velocity of native (full-length) myosin XI. The estimated velocities of native (full-length) *Cb*XI-1 and *Cb*XI-2 were 73 μm s^−1^ (14.5 μm s^−1^ × 5) and 66 μm s^−1^ (13.2 μm s^−1^ × 5), respectively (Fig. 3 and Table 1). The estimated velocities of *Cb*XI-1 and *Cb*XI-2 are therefore almost the same as the cytoplasmic streaming velocity in members of the genus *Chara* (*C. corallina* and *C. braunii*), which have velocities of 70 μm s^−1^ (13, 17).

**Table 1.** ***V***_max_ and ****K****_app_ of actin-activated ATPase activity and actin-sliding velocity of *Chara* myosin^1^.

|  | *CbXI-1* | *CbXI-2* | *CbXI-3* | *CbXI-4* | *CcXI*<sup>2</sup> |
| --- | --- | --- | --- | --- | --- |
| $V_{\max}$ (Pi s<sup>-1</sup> head<sup>-1</sup>) | 410 | 200 | 260 | 230 | 580 |
| $K_{\text{app}}$ (μM) | 46 | 45 | 15 | 8.7 | 23 |
| Velocity of MD<br>(μm s<sup>-1</sup>) | 15 ± 0.7 | 13 ± 0.6 | 3.0 ± 0.2 | 3.1 ± 0.3 | 4.7 ± 0.3 |
| Velocity of full-length myosin<br>expected from MD<sup>3</sup> (μm s<sup>-1</sup>) | 73 | 66 | 15 | 16 | 24 |
| Velocity of chimeric full-length<br>myosin<sup>4</sup> (μm s<sup>-1</sup>) | 50 ± 3.7 | ND | ND | ND | 16 ± 0.9 |
| Duty ratio<sup>5</sup> (%) | 9.6 | 5.4 | 30 | 26 | 43 |

<sup>1</sup> Measured at 25 °C
<sup>2</sup> $V_{\max}$ , $K_{\text{app}}$ , and velocity of *CcXI* MD (23), velocity of chimeric full-length *CcXI* (10)
<sup>3</sup> Calculated by the method described in (33)
<sup>4</sup> Chimeric myosins were made by replacing six IQ motifs of *Chara* myosins with those of *AtXI-2*
<sup>5</sup> Calculated by the $V_{\max}$ values of the actin-activated ATPase activities and the velocities of MD as described in (33)

**Figure 3.**
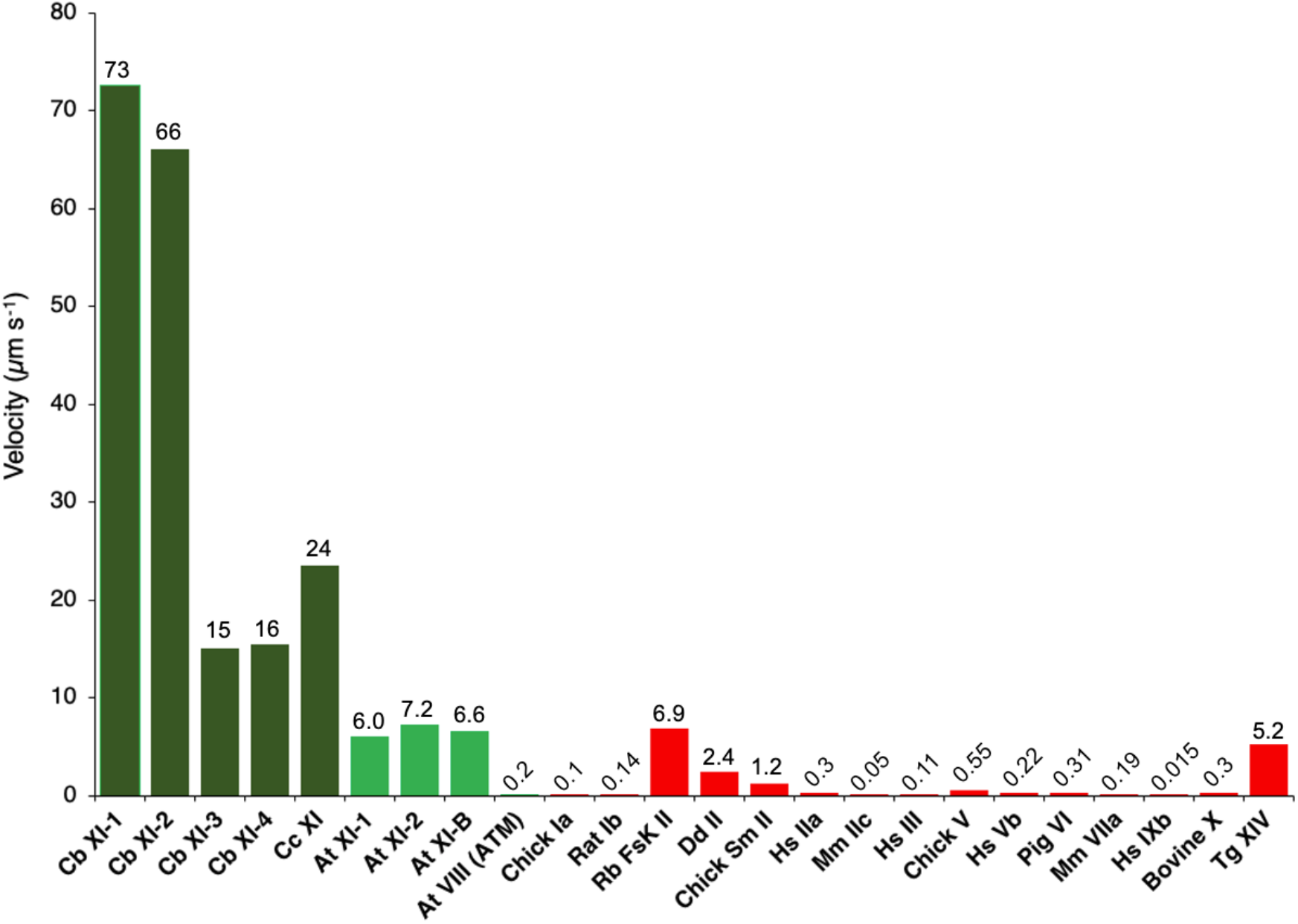
Velocities of various classes of myosins. *Cb*XI: *Chara braunii* myosin XI (this paper), *Cc*XI: *Chara corallina* myosin XI (22, 23), *AtXI*: *Arabidopsis thaliana* myosin XI (10, 33), *At*VIII: *Arabidopsis thaliana* myosin VIII (4), Chick Ia: Chicken myosin Ia (44), Rat Ib: Rat myosin Ib (45), Rb FskII: Rabbit fast skeletal myosin II (46), *Dd* II: *Dictyostelium discoideum* myosin II (47), Chick Sm II: Chicken gizzard smooth muscle myosin II (48), *Hs* IIa: *Homo sapiens* myosin IIa (49), Mm IIc, *Mus musculus* myosin IIc (50), *Hs* III: *Homo sapiens* myosin III (51), Chick V: Chicken myosin Va (52), *Hs* Vb: *Homo sapiens* myosin Vb (53), Pig VI: Pig myosin VI (54), Mm VIIa: *Mus musculus* myosin VIIa (55), Hs IXb: Homo sapiens myosin IXb (56), Bovine X: Bovine myosin X (57), Tg XIV: *Toxoplasma gondii* myosin XIV (58)

To further estimate the velocity of *Cb*XI-1, we used a chimeric *Cb*XI-1 with the same lever arm length as native *Cb*XI-1 (chimeric *Cb*XI). The velocity of the chimeric *Cb*XI-1 was 50 ± 3.7 μm s^−1^ (Table 1). This velocity is somewhat less than the velocity of *Cb*XI-1 estimated from *Cb*XI-1 MD. The smaller velocity of the chimeric *Cb*XI-1 may be due to improper linkage between *Cb*XI-1MD and the IQ motifs of *At*XI-2 or to improper interactions between *Cb*XI-1MD and myosin light chains (calmodulin) of *At*XI-2. This velocity of the chimeric *Cb*XI-1 (50 μm s^−1^) was three times faster than the velocity of the chimeric *Cc*XI (16 μm s^−1^) (10), which had the same IQ motifs and the same coiled-coil and tail domains as the chimeric *Cb*XI-1 (both chimeric *Chara* myosins had IQ motifs, coiled-coil, and tail domains of *Arabidopsis* myosin XI-2). These results strongly suggest that *Cb*XI-1 and *Cb*XI-2 are the myosins causing cytoplasmic streaming in *Chara braunii* and demonstrate that *Cb*XI-1 is the fastest myosin yet measured among all organisms (Fig. 3).

### ATPase activities

The ATPase activities of *Cb*XI-1, *Cb*XI-2, *Cb*XI-3, and *Cb*XI-4 MDs were plotted as a function of actin concentration and fit to the Michaels–Menten equation to determine the maximum rate of ATP turnover (***V***_max_) and the actin concentration at which the ATPase rates were one-half of the maximal rate (***K***_app_). The ***V***_max_ and ***K***_app_ values of *Cb*XI-1 MD were 410 Pi s^−1^ head^−1^ and 46 μM, respectively (Table 1). This ***V*_max_** value is similar to that of *Cc*XI MD (21, 22), although the actin velocity of *Cb*XI-1 MD is three times faster than that of *Cc*XI MD. The ***V*_max_** of actin-activated ATPase activities was not correlated with actin velocity among the four *Cb*XIs. This discrepancy may arise because the rate-limiting step of ***V*_max_** of actin-activated ATPase activities (phosphate dissociation from actin–myosin–ADP·Pi complex) and the rate-limiting step of actin velocity (ADP dissociation from actin–myosin–ADP complex) are different (22).

### Crystal structure of the myosin XI MD

To investigate the molecular mechanism of the ultrafast movement of *Cb*XI-1, we tried to crystallize *Cb*XI-1 MD. However, we were unsuccessful, probably because *Cb*XI-1 MD is unstable and prone to semi-denaturation. ATPase activity of *Cb*XI-1 tends to drop in a relatively short time compared to other myosins, which makes it difficult to obtain the crystal structure. Such instability of ATPase activity was observed for all *Chara* myosins. Although class XI myosin is the fastest myosin class in the myosin superfamily (Fig. 3), the atomic structure of the class XI myosin MD has yet to be solved. Therefore, in this study, we decided to clarify the structural features responsible for the high velocity of class XI myosins by crystallographic analysis of other class XI myosin. We chose *Arabidopsis* myosin MYA2 (*At*XI-2) as the crystallization target, because *At*XI-2 has a standard velocity among class XI myosins, it is faster than most animal myosins (Fig. 3) (33), and its ATPase activity is more stable than those of *Chara* myosins. *At*XI-2 has an amino acid sequence relatively similar to that of *Cb*XI-1 MD, having 63% identity and 87% similarity.

We succeeded in solving the crystal structure of *At*XI-2 MD bound with ADP and AlF_4_ ^−^ at 2.8 Å resolution, which is the first atomic structure of the class XI myosin MD (Fig. 4 *A* and *B*, and Supplementary Table 1). Figure 4 *C* and *D* show comparisons of the structure of ADP·AlF_4_ ^−^-bound *At*XI-2 with the structures of ADP·AlF_4_ ^−^-bound other classes of myosins. Although there are some deviations in the position of the main chain, the nucleotide interaction region is almost identical (Fig. 4 *C*). The backdoor size for phosphate dissociation of *At*XI-2 was also very similar to those of myosins of other classes (Fig. 4 *D*). Thus, the structure near the nucleotide-binding region of myosin XI was not markedly different from those of myosins of other classes (myosin II, myosin VI, and myosin XIV).

**Figure 4.**
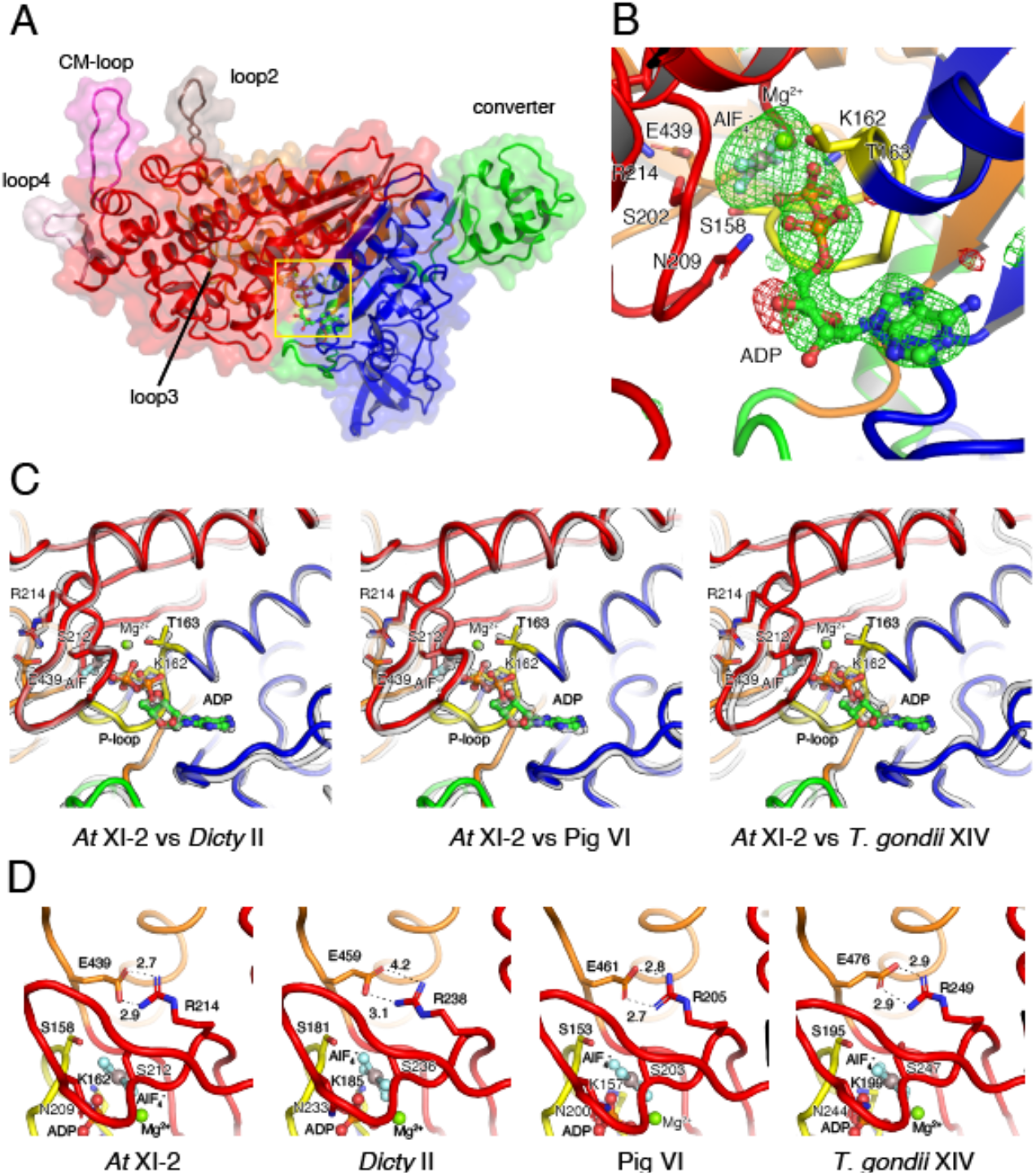
Crystal structure of *At*XI-2 MD bound with ADP and AlF_4_^−^. (*A*) Main surface loops (loops 1, 2, 3, and 4 and the CM-loop) of *At*XI-2 MD are shown. Upper 50k, lower 50k, N-terminal, and converter subdomains are colored in red, orange, blue, and green, respectively. (*B*) The nucleotide-binding region of *At*XI-2 MD. The |Fo|-|Fc| map calculated without ADP:Mg^2+^ and AlF_4_ at the binding pocket contoured at 4.0 sigma are shown in red (negative) and green (positive). (*C*) Comparison of the nucleotide-binding regions in various myosins (gray) bound to ADP and AlF_4_^−−^. (*D*) Comparison of the size of the backdoor for the Pi release in various myosins bound to ADP and AlF_4_ ^−^. *At*XI-2, *Dictyostelium* myosin II (PDB: 1MND), Pig myosin VI (PDB: 4ANJ), *Toxoplasma gondii* myosin XIV (PDB: 6DUE).

Myosins of different classes have significantly different motor properties, such as velocity, ATPase activity, and duty ratio. Even within the same class, the motor properties are different for each myosin. The differences in the properties of each myosin motor are due to differences in the MD’s amino acid sequence. Therefore, we investigated the structural regions in the MD with large variations in amino acid sequence among various myosin classes (Fig. 5). A comparison of the amino acids of different myosins showed that the central part of the MD is highly conserved, and these regions are responsible for basic chemical-mechanical energy conversion. In addition to the src-homology 3 region of the N-terminal subdomain, the actin-binding regions of the upper 50k and lower 50k subdomains have high amino acid diversity among various myosin classes (Fig. 5 *B*: Various classes and *SI Appendix*, Fig. S1). The same tendency was observed among myosin XIs, although weaker than among various myosin classes. (Fig. 5 *B*: XIs). These regions would be responsible for the diversity of motor properties among myosins.

**Figure 5.**
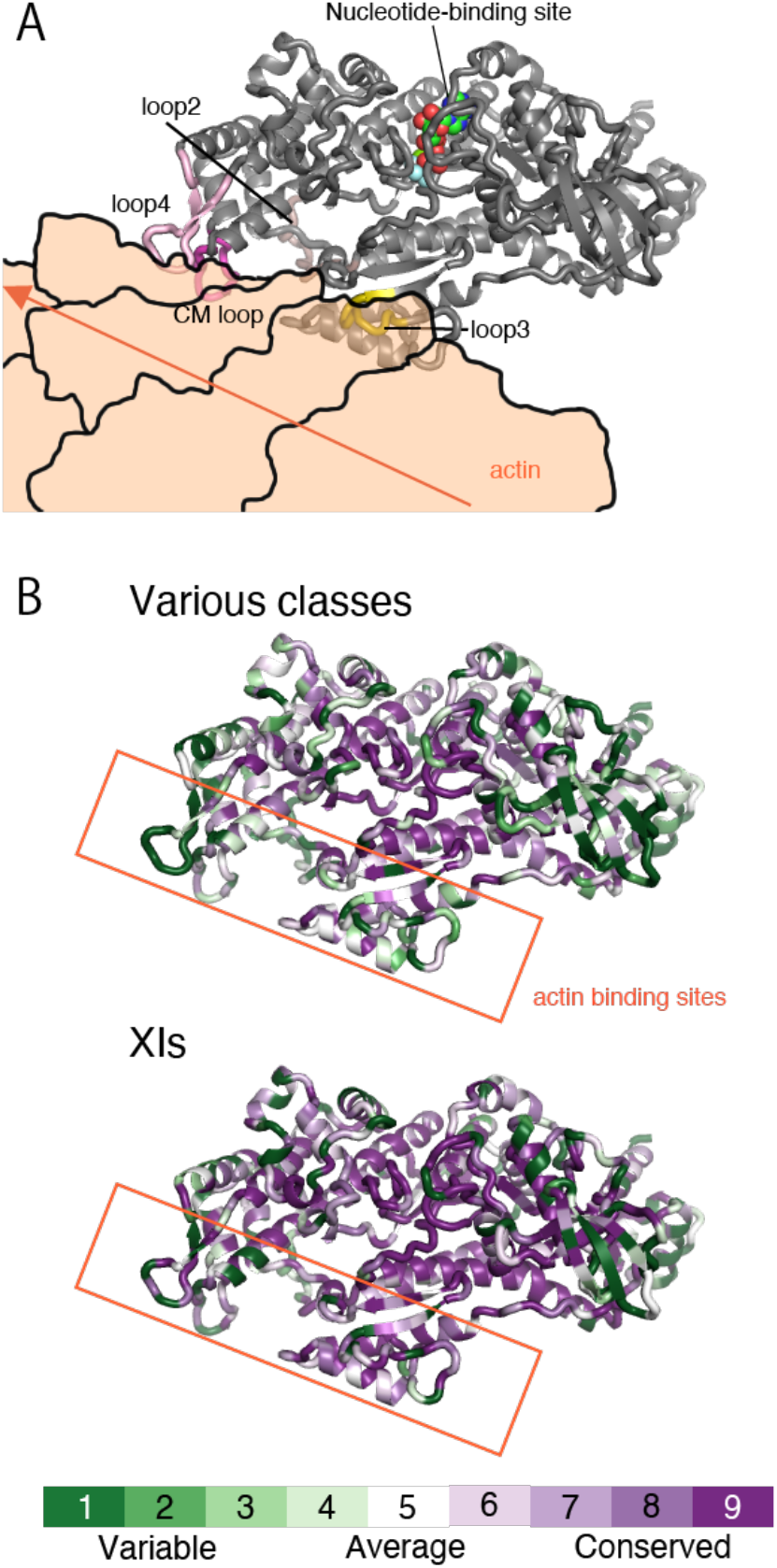
Actin-binding region with high amino acid diversity among myosins. (*A*) Docking model of *A*tXI-2 MD and actin. This model was created by replacing myosin X in 5KG8 (Rigor myosin X co-complex with an actin filament) with *At*XI-2 MD. The actin in 5KG8 was replaced by 6BNO (structure of bare actin filament). (*B*) Heat map visualization of *A*tXI-2 MD showing amino acid conservation and diversity, generated using ConSurf (http://consurf.tau.ac.il/). The conservation score is calculated using the Maximum Likelihood paradigm. The amino acids of myosins are colored by conservation score ranging from green (1; most variable) to purple (9; most conserved residues), as shown in the color legend. The rate was changed to 1 for residues with four or fewer sequence overlaps in the alignment. **Various classes** showing amino acid comparison of 10 classes of myosins: class I (human Ic and Rat Ib), class II (*Dictyostelium* II, chicken fast skeletal muscle, chicken smooth muscle, and rabbit skeletal muscle), class V (chicken Va and human Vb), class VI (pig VI), class VII (*Drosophila* VIIa and mouse VIIb), class VIII (*Arabidopsis* VIIIa and *Arabidopsis* VIIIb), class IX (human IXa and human IXb), class X (cow X), class XI (*Arabidopsis* XI-2, *Chara corallina* XI, and *Chara braunii* XI-1), and class XIV (*Toxoplasma* XIV). **XIs** showing the amino acid comparison of 18 myosins belonging to class XI: *Arabidopsis* XI-1, *Arabidopsis* XI-2, *Arabidopsis* XI-A, *Arabidopsis* XI-B, *Arabidopsis* XI-C, *Arabidopsis* XI-D, *Arabidopsis* XI-E, *Arabidopsis* XI-F, *Arabidopsis* XI-G, *Arabidopsis* XI-H, *Arabidopsis* XI-I, *Arabidopsis* XI-J, *Arabidopsis* XI-K, *Chara corallina* XI, *Chara braunii* XI-1, *Chara braunii* XI-2, *Chara braunii* XI-3, and *Chara braunii* XI-4).

### Velocities of actin-binding surface loop mutants

Since the actin-binding loops 2 and 3 contribute to the high velocity of *Cc*XI (23), these loops may be involved in the difference in velocity between *Chara* myosin subgroups 1 and 2 (Tables 1 and 2). To investigate this possibility, we made mutated *Cc*XI constructs in which the *Cc*XI actin-binding loop 2, loop 3, or both were replaced by those of *Cb*XI-1. The loop 3 substitution mutations had little effect on the velocity. However, replacing loop 2 of *Cc*XI with that of *Cb*XI-1 increased the velocity of the mutated *Cc*XI by a factor of 1.3 to 1.4 (Table 3). We have previously shown that when two or more consecutive positive charges are added to loop 2, and the net charge increases by more than two, the velocity of *Cc*XI decreases (23). In the mutation in which loop 2 of *Cc*XI was replaced with that of *Cb*XI-1, there was no continuous addition of positive charge, and the net charge of loop 2 increased by only one. This change in loop 2 may reduce the velocity slightly but is unlikely to increase the velocity. Loop 2 of *Cb*XIs (subgroup 1) is characterized by three consecutive proline and five glycine clusters (Table 2). Proline and glycine clusters are known to disrupt the secondary structure (34, 35), so a *Cb*XI-1 loop 2 with this feature would be a completely free loop structure, resulting in increased flexibility. The increased flexibility of loop 2 may allow for fast ADP dissociation by facilitating the transition to the strong-binding (rigor) state.

**Table 2.**
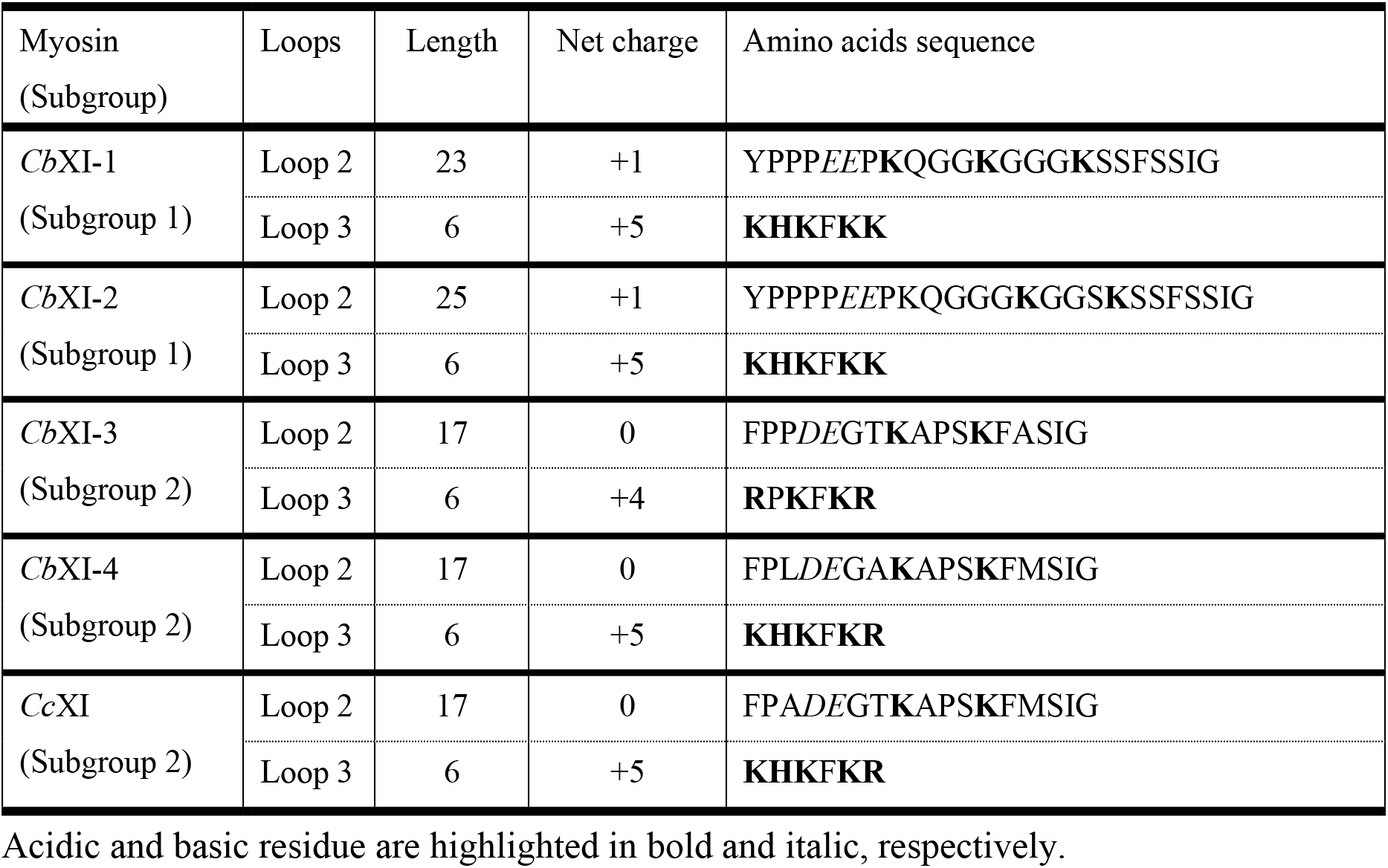
Loop 2 and loop 3 sequences of *Chara* myosins and velocities of their MD constructs.

## Discussion

In this work, we successfully cloned the fastest myosin in the biological world, *Cb*XI-1. We also succeeded, for the first time, in solving the atomic crystal structure of the MD of class XI myosin. The amino acid sequence features of *Cb*XI-1, mutation experiments, and the crystal structure of class XI myosin indicate that the actin-binding sites are crucial for defining the myosin velocity.

The myosin superfamily currently has 79 classes, each containing several subclasses (1). Of the 79 myosin classes, 17 classes are found in animals, and 2 are found in green plants. Other myosin classes are found in fungi and Protista. The velocities and ATPase activities of most of the 19 myosin classes present in vertebrates and plants have been measured. The velocities of myosins differ greatly depending on the class and subclass. Although skeletal muscle myosin II is exceptionally fast, most animal myosin has a velocity of less than 1 μm s^−1^. In contrast, plant-specific class XI myosins have velocities of 5 μm s^−1^ or higher (Fig. 3). The high velocity of class XI myosins is thought to be related to the plant-specific role; the main role of class XI myosins is to drive cytoplasmic streaming, whereas the role of most animal myosins is the generation of force. The force generated by class XI myosin is smaller than that produced by animal myosins (36). Because plant cytoplasmic streaming is caused by myosin XI-bound organelle movement along actin filaments, the velocity of cytoplasmic streaming in plants is thought to be approximately equal to the velocity of the class XI myosin of that plant. Based on the cytoplasmic streaming velocity in the aquatic genus *Chara*, it has long been proposed that *Chara* possesses a myosin with a velocity of 70 μm s^−1^ (13–17), but the myosin gene responsible for this has not yet been identified. This work revealed that the myosin gene is *Cb*XI-1 and *Cb*XI-2.

The *Chara* myosin XIs have expanded independently of the expansion of the land plants (37), resulting in four genes. *Cb*XI-1 and *Cb*XI-2, the members of subgroup 1, form a clade having a longer branch than that of a clade formed by *Cb*XI-3, *Cb*XI-4, and *Cc*XI, which are in subgroup 2 (Fig. 2). Because *Cb*XI-4 and *Cc*XI form a clade, they are putative orthologs. The longer branch lengths in the *Cb*XI-1 and *Cb*XI-2 clade imply the action of adaptive evolution to increase the velocity, as they have fastest movement and do not seem to have lost any functionality. The genome project found that *Cb*XI-1 is the most highly expressed myosin XI in the whole plant (*SI Appendix*, Table S2) (26). Thus, *Cb*XI-1 appears to be the main contributor to cytoplasmic streaming in *Chara braunii*. *Cb*XI-2, which has almost the same velocity as *Cb*XI-1, would provide redundancy of the function of *Cb*XI-1.

We estimated the duty ratio of *Cb*XI myosins from the ****V****_max_ values of the actinactivated ATPase activities and the velocities of MD (Table 1). The duty ratios of subgroup 2 myosins are much higher than those of subgroup 1. In order for a myosin-bound vesicle to remain associated with actin filaments, and to move continuously along the actin filaments, at least one of the myosin MDs on the vesicle must always be in strongly bound to the actin filaments. Therefore, the reciprocal of the duty ratio is the lowest number of MDs on a vesicle required for continuous movements of the myosin-bound vesicle on the actin filaments. Subgroup 2 myosins, which have a high duty ratio, can transport vesicles with fewer myosin MDs than can subgroup 1 myosins. Subgroup 2 myosins may function in small vesicle transport, in which the number of bound myosins is limited.

We have obtained the first crystal structure of class XI myosin at 2.8 Å resolution, which is the fastest of all the measured myosin classes. The structure of class XI myosin, *At*XI-2 near the nucleotide-binding region was similar to those of other classes of myosins: *Dictyostelium* myosin II, pig myosin VI, and *Toxoplasma* myosin XIV (Fig. 4 *C* and *D*). This finding is consistent with a previous report that the central part of the myosin structure is similar among different classes of myosins, even though their velocities and ATPases are very different (38). However, there is considerable diversity among the amino acids in the actin-binding sites of myosins (Fig. 5). Most recently, the rigor structure (post-power stroke, non-nucleotide state) of acto-*Cc*XI determined by cryo-EM at 4.3 Å resolution was reported (25). The cryo-EM and our crystal structures show that class XI and other classes of myosins (39–41) differ in their actin-binding mode (*SI Appendix*, Fig. S1). These results suggest that actin-binding sites determine the diversity of velocities among the classes and subclasses of myosins. Kinetic data from various classes of myosin also support this suggestion. The dissociation rate of ADP from the myosin–ADP complex in the absence of actin is nearly identical among myosins with different velocities. In contrast, the ADP dissociation rate from the actin–myosin–ADP complex, which is the rate-limiting step in myosin velocity, differs depending on the classes and subclasses of myosins (22). Different classes and subclasses of myosins bind to actin in somewhat different ways, which probably cause different changes in the nucleotide-binding site through communication between the actin- and the nucleotide-binging sites within the MD, resulting in different kinetics. The recently reported cryo-EM structure of acto-*Cc*XI MD was in the non-nucleotide state (25). When the cryo-EM structure of acto-*Chara* MD bound to ADP is revealed, we would know the structural reason why the ADP dissociation rate from acto-*Chara* MD-ADP, i.e., the velocity of *Chara* myosin, is so fast.

We have previously shown that the actin-binding loops 2 and 3 contribute to the high velocity of *Cc*XI (23). The differences in the amino acid sequence of loop 2 between subgroup 1 and subgroup 2 of *Chara* myosins is marked (Table 2). We made mutated *Cc*XI (subgroup 2) constructs that replace the *Cc*XI loop 2, loop 3, or both with those of *Cb*XI-1 (subgroup 1). Mutated *Cc*XI constructs in which loop 2 was replaced with that of *Cb*XI-1 increased the velocity by 1.3–1.4 times (Table 3). Because this mutation increases the net positive charge of loop 2 of *Cc*XI by only one, and this small positive charge increase of loop 2 would not change or slightly reduce the velocity of *Cc*XI (23), this increase in velocity may not be due to the increase in net charge of loop 2. Loop 2 of *Cb*XI-1 (subgroup 1) is characterized by proline and glycine clusters (Table 2), which are known to disrupt the secondary structure (34, 35). Loop 2 of *Cb*XI-1 (subgroup 1) with this feature is probably a complete loop and is highly flexible. Myosin binds to actin through many regions in the upper 50k and lower 50k subdomains. The flexibility of loop 2 of *Cb*XI-1 may facilitate strong binding to actin in regions other than loop 2, which would promote rapid ADP dissociation. Because the difference in velocity between *Cc*XI and *Cb*XI-1 is about three times, and loop 2 of *Cb*XI-1 contributes only half of this difference, other actin-binding sites could contribute to the difference in velocity. Actin-binding sites in myosins are large regions containing upper 50k and lower 50k domains, and many of these domains have tertiary structures. Since mutations in these tertiary structures can disrupt the overall structure, mutation experiments on these structures would be difficult.

**Table 3.**
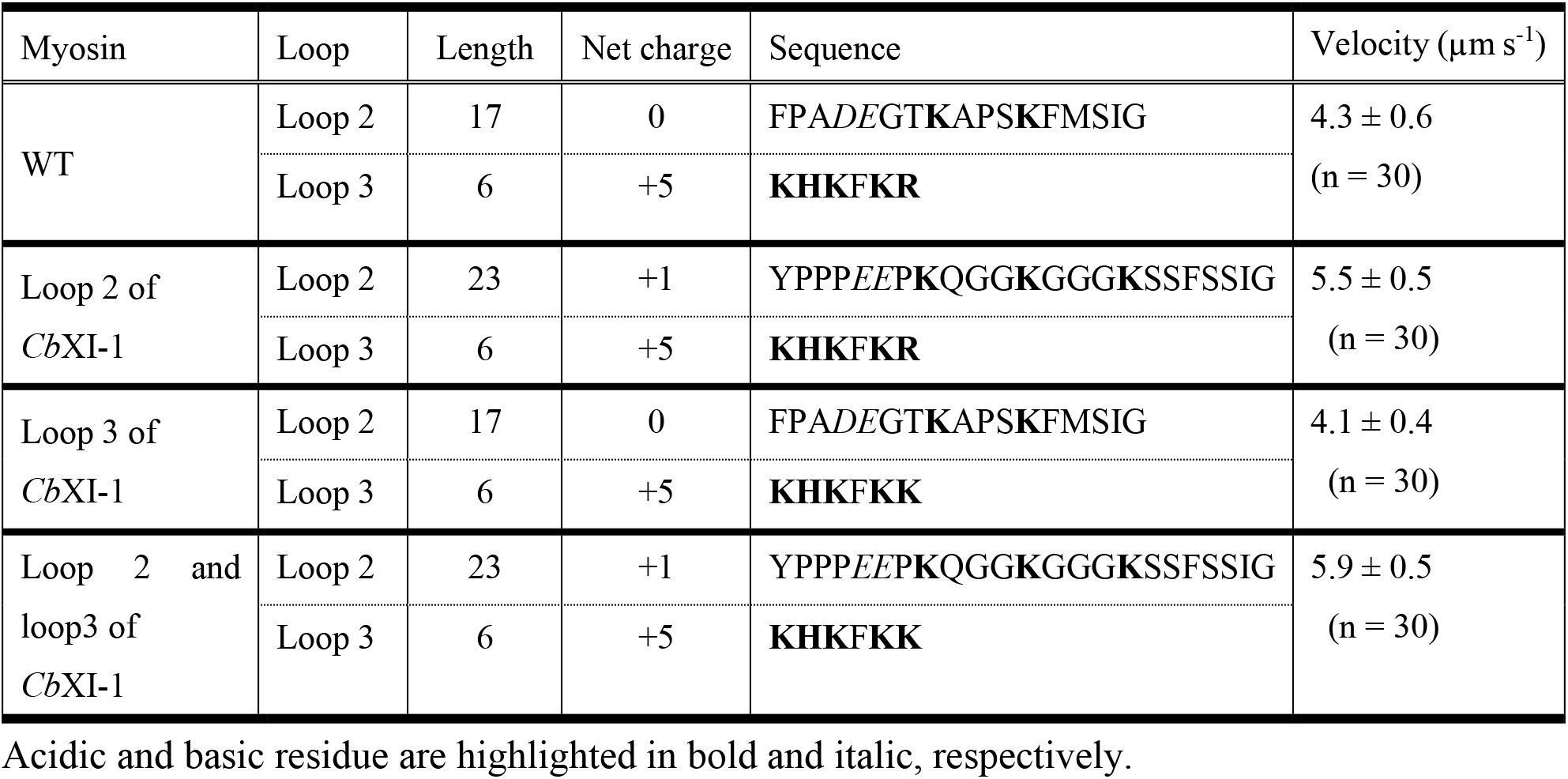
Amino acid sequence and actin sliding velocity of WT and loop 2, -loop 3 and -loop 2, 3 mutants of *Cc*XI.

Detailed kinetics analyses and single molecule assays using the fastest myosin, *Cb*XI-1, would provide further insights into the detailed chemical-mechanical conversion mechanism of myosin. The gene for *Cc*XI, the fast myosin, has contributed to the development of nano-machines (24, 25) and the enhancement of plant growth (10, 42). The gene for *Cb*XI-1, the ultrafast myosin, will greatly contribute to a variety of research and development.

## Materials and methods

described in *SI Appendix, Materials and Methods*

## Data Availability

The reconstructed nucleotide sequence of *Cb*XI-4 mRNAs are deposited in DDBJ under accession nos. BR001749 and BR001750.

The atomic coordinates and structure factors of *At*XI-2 MD have been deposited in the Protein Data Bank under the accession code 7DHW (http://dx.doi.org/10.2210/pdb7DHW/pdb).

## Acknowledgements

The synchrotron radiation experiments were performed at Photon Factory (proposals 2016G048, 2016R-19). We thank the beamline staff at BL1A and BL17A of Photon Factory (Tsukuba, Japan) for help during data collection. We also thank Ms. Mari Udo for color coordinate of Fig.1 and Enago (www.enago.jp) for the English language review. This work was supported by Grant-in-Aid for Scientific Research (Grant Numbers JP 20001009 to M.T, JP 25221103 to M.T, JP 15K07185 to H.S., JP 16H05764 to H.S., JP 18K06382 to H.S., 15H04413 to T.N, JP 18H05425 to T.M., JP 20K06583 to K.I., JP 17K07436 to K.I., JP 17K07436 to K.I., 15H01309 to K.I.) from Japan Society for the Promotion of Science (JSPS), by ALCA (JPMJAL1401 to M.T. and KI) from the Japan Science and Technology Agency (JST), and by Basis for Supporting Innovative Drug Discovery and Life Science Research (BINDS; JP20am0101083 to T.M.) from Japan Agency for Medical Research and Development (AMED).

## Supplementary Information

### Materials and Methods

#### RNA extraction

Thalli of strain S276 were harvested in soil-water medium for the Charales (SWC-3) (1) under controlled laboratory conditions at 23ºC with a 16-h light: 8-h dark cycle with 24.5 μmol photons m s illumination provided by fluorescent lamps. The upper parts of thalli with reproductive organs were collected, frozen in liquid nitrogen, and stored at −80ºC until further processing. Frozen samples were ground in liquid nitrogen. Total RNAs were then extracted with ISOGEN (Nippon Gene, Tokyo, Japan), and purified using the QIAGEN RNeasy Plant Mini Kit.

#### Cloning of motor domain of *Chara braunii* myosins

cDNA of *Cb*XI-1 (g50407), *Cb*XI-2 (g48390), *Cb*XI-3 (g24025), and *Cb*XI-4 (g48658) motor domains (MDs) were amplified from total RNA of *Chara braunii*. The primers sequences used are as follows: 5’-TAGCGTCTCTTCAAAATGGCA-3’ and 5’ CGCACTCTCCTTGTCATCTTCTT-3’ for *Cb*XI-1, 5’ TATTTATAGTTCAGAATGGCGGAGC-3’ and 5’-CCTGCGGCCAATTCTTTT-3’ for *Cb*XI-2, 5’-CTCAGGAGTGTCACCATGGG-3’ and 5’ ACTCTGCGCTGGATCTTGAC-3’ for *Cb*XI-3, 5’ CTCACTCAGAATCATCATGGGGTC-3’ and 5’ CTTCATTCTCTCATAGTCTTTACGCATCAG-3’ for *Cb*XI-4.

#### Protein engineering, Expression, and Purification

##### CbXI-1, *Cb*XI-2, *Cb*XI-3, *Cb*XI-4 MDs and chimeric *Cb*XI-1

A baculovirus transfer vector for *Cb*XI-1, *Cb*XI-2, *Cb*XI-3, *Cb*XI-4 MDs were generated as follows: The cDNAs of motor domains of *Cb*XI-1 (amino acid residues 1– 745 of *Cb*XI-1), *Cb*XI-2 (amino acid residues 1–751 of *Cb*XI-2), *Cb*XI-3 (amino acid residues 1–747 of *Cb*XI-3), *Cb*XI-4 (amino acid residues 1–742 of *Cb*XI-4) with optimized insect *Trichoplusia ni* codons were artificially synthesized by Eurofins Genomics because the expression levels in insect cells were very low when native cDNAs were used. These were cloned into the pFastBac MD (2) using the In-fusion cloning kit (Takara). The resulting constructs, pFastBac *Cb*XI-1, *Cb*XI-2, *Cb*XI-3, *Cb*XI-4 MDs encode an N-terminal amino acids (MDYKDDDDKRS) containing the FLAG tag (DYKDDDDK), amino acid residues 1–745, 1–751, 1–747 or 1–742 of *Cb*XI-1, *Cb*XI-2, *Cb*XI-3, *Cb*XI-4, respectively, C-terminal amino acids (GGGEQKLISEEDLHHHHHHHH SRMDEKTTGWRGGHVVEGLAGELEQLRARLEHHPQGQREPSR) containing a flexible linker (GGG), a Myc-epitope sequence (EQKLISEEDL), His tag (HHHHHHHH) and SBP tag (MDEKTTGWRGGHVVEGLAGELEQLRARLEHHPQGQREP).

A baculovirus transfer vector for chimeric *Cb*XI-1 was generated as follows: cDNA of *Cb*XI-1 was mutated to create annealing sites for pFastBac *At*XI-2 neck-tail (3). It was ligated to pFastBac *At*XI-2 neck-tail using the In-Fusion cloning kit (Takara). The resulting construct pFastBac chimeric *Cb*XI-1 encodes N-terminal amino acids (MDYKDDDDKRS) containing the FLAG tag, amino acid residues 1–741 of *Cb*XI-1, amino acid residues 735–1,505 of *At*XI-2, C-terminal amino acids (GGGEQKLISEEDLHHHHHHHHSRMDEKTTGWRGGHVVEGLAGELEQLRARL EHHPQGQREPSR) containing a flexible linker, a Myc-epitope sequence, His tag and SBP tag.

Mutated *Cc*XI MDs with the amino acid sequence of the loop 2 and/or loop 3 regions changed to those of *Cb*XI-I.

Eurofins Genomics made mutational *Cc*XI MD constructs in which loop 2 and/or loop 3 of *Cc*XI MD were changed to those of *Cb*XI-1.

These constructs were expressed in insect cells (High Five, Life Technologies) and purified using Ni-affinity and FLAG-affinity resins as previously described (4).

##### *At*XI-2 MD

A baculovirus vector for *At*XI-2 MD motor domain (pFastBac Flag His TEV *At*XI-2 MD) was generated as follows: The motor domain of *At*XI-2 myosin contains residues 1-738 of the *At*XI-2 myosin heavy chain. The motor domain of *At*XI-2 was amplified by PCR. The gel-purified PCR fragment was cloned into the pFastBac vector using the In-fusion cloning kit (Takara). The resulting protein, *At*XI-2 MD has N-terminal amino acids (MDYKDDDDKRSMHHHHHHDYDIPTTENLYFQGA) containing the FLAG tag, His tag and TEV (ENLYFQG), amino acid residues 1-738 of *At*XI-2. *At*XI-2 MD was expressed in insect cells (High Five, Life Technologies) and purified using nickel-affinity and FLAG-affinity resins as previously described (4).

#### ATPase activity

ATPase activities were determined by measuring released phosphate as previously described (5). The reaction mixtures for the assay of actin-activated Mg^2²^-ATPase activity contained were done in 25 mM KCl, 4 mM MgCl_2_, 25 mM Hepes-KOH (pH 7.4), 2 mM ATP, 1 mM DTT, and 1 mg/ml BSA and at 25ºC, 0.125 4 mg/ml F-actin.

#### *in vitro* gliding assay

The velocity was measured using an anti-myc antibody-based version of the *in vitro* actin filament gliding assay as previously described (2). The velocity of actin filaments was measured in 150 mM KCl, 4 mM MgCl_2_, 25 mM Hepes-KOH (pH 7.4), 2 mM ATP, 10 mM DTT and oxygen scavenger system (120 μg/ml glucose oxidase, 12.8 mM glucose, and 20 μg/ml catalase) at 25ºC. Average sliding velocities determined by measuring the displacements of actin filaments.

#### Crystallization and Data Collection

Purified sample was incubated with 200 μM ADP, 1 mM AlCl_3_, and 5 mM NaF in this order for every 30 min on ice. The protein solution (0.1 μl; 4 mg/ml) was mixed with the reservoir solution (0.1 μl) consisting of 0.1 M MMT buffer (pH 6.0; Molecular Dimensions) and 25% PEG-1500. Crystals had grown at 296 K in sitting drops by vapor diffusion. The crystals were soaked for 1 min in 0.1 M MMT buffer (pH 6.0), 25% PEG 1500, 200 μM ADP, 1 mM AlCl_3_, and 5 mM NaF and 20% glycerol, and were frozen and stored in liquid nitrogen.

The X-ray diffraction data were collected from a single crystal at a cryogenic temperature (100 K) on BL-17A at the Photon Factory (Tsukuba, Japan). The collected data were processed using XDS software (6). The structure was solved by molecular replacement with Phaser (7) as a search model for *Dictyostelium Discoideum* MD complexes with ADP-AlF_4_ (PDB ID code; 1MND). The atomic model was built using Coot (8) and iteratively refined using Phenix (9). TLS (Translation/Libration/Screw) refinement was performed in late stages of refinement. The refined structures were validated with RAMPAGE (10). All molecular graphics were prepared using PyMOL (The PyMOL Molecular Graphics System, Version 2.1.1, Schrodinger, LLC, New York, NY, USA).

#### The phylogenetic tree of *Chara* myosin XIs and the whole of myosin XIs

Myosin homologs were collected from NCBI refseq database for *Arabidopsis thaliana*, *Amborella trichopoda*, and *Selaginella moellendorffii*, and *Physcomitrium patens* based on BLASTP search using the *Cb*XI-1 as a query. Another search for NCBI nr dataset were performed with organisms specified as Viridiplantae but excluding Spermatophyta. Further, ferns (*Salvinia cucullata* and *Azolla filiculoides*) (11), bryophytes (*Marchantia polymorpha* (12) and *Anthoceros agrestis* (13)), and Zignematophycean algae (*Spirogloea muscicola* and *Mesotaenium endlicherianum*) (14) were searched for respective datasets. Those datasets were obtained from ftp://ftp.fernbase.org/Salvinia_cucullata/Salvinia_asm_v1.2, ftp://ftp.fernbase.org/Azolla_filiculoides/Azolla_asm_v1.1, https://phytozome.jgi.doe.gov/pz/portal.html#!info?alias=Org_Mpolymorpha, https://www.hornworts.uzh.ch/static/download/a_agr_oxford.zip, and https://figshare.com/articles/Genomes_of_subaerial_Zygnematophyceae_provide_insights_into_land_plant_evolution/9911876/1, respectively.

Thus, 80 sequences were retrieved and aligned with einsi in MAFFT v7.475 (15). The alignment were read with Mesquite version 3.61 (16) and well-aligned regions were marked as included after excluding all characters, to ensure only well-aligned regions were used for the subsequent analyses. Two sets of matrix were extracted from the entire alignment; one including myosin VIII, and another confined to myosin XI excluding sequences having deletions in conserved regions, which may be either alternative splicing or misannotation. Sequences identical in the included region were regarded as a single OTU and one representative was retained during the analysis. Thus, green plant myosin dataset included 567 sites from 43 OTU and myosin XI dataset included 1060 sites from 32 OTU. Substitution model were chosen using ProteinModelSelection.pl from RAxML distribution. Standard RAxML version 8.2.12 (17) was used for the maximum likelihood tree search with -f a -# 100 option and 24 bits of random seed were provided from system random source /dev/urandom to -p and -x options. 1000 bootstrap dataset were prepared using seqboot from PHYLIP version 3.697 (18) and split to individual dataset and then run essentially the same RAxML command. The bootstrap trees were collected along with the original tree amplified 1000 times to force the consensus topology and the bootstrap value were counted using consense then subtracted for 1000 to recover the bootstrap value.

#### Identification of full-length sequence of *Cb*XI-4

Initially, g48658 was identified as a Myosin XI homolog and designated *Cb*XI-4. The g48658 spanned almost entire range of the scaffold_743 and appeared to contain only N terminal part. Thus, the transcript sequence (2554 bp) was used as a query to search several transcriptome assemblies of *Chara braunii*, which were constructed during the genome project but was not described (ref genome paper). A BLASTN (19) search to an assembly constructed with TransAbyss (20) found a 4242-bp contig, which is considerably longer than the query. This sequence was used to search the predicted transcriptome and found that g66754 have partially matching sequence without scattered mismatches. g66754 was located on the minus strand at the 3’ end of scaffold_8 and RNA-seq data suggested transcripts spanned to g66753 and g66745. The prediction by augustus (21) contained exons that was not linked with RNA-seq data overlapping introns and having similarity to retrotransposons. Thus, the g66745-g66753-g66754 locus was manually edited on WebApollo (22) to conform with the RNA-seq data with intron support, and two isoforms were retained. This sequence was connected to the 4242-bp contig of the RNA-seq assembly to end up in the two full-length nucleotide sequence of the transcripts.

### Supplementary Text

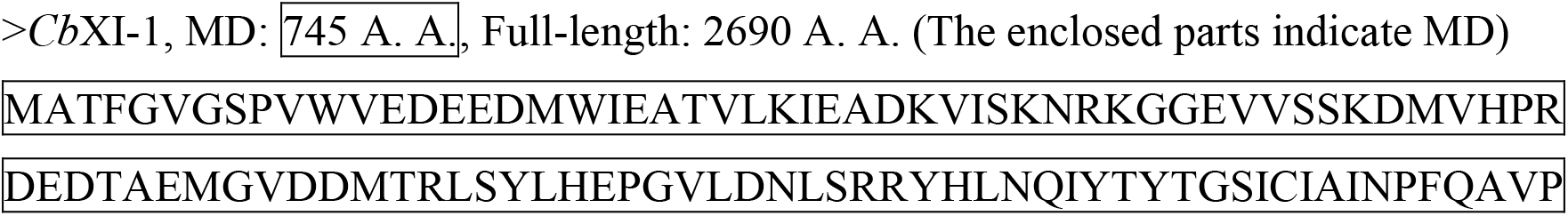
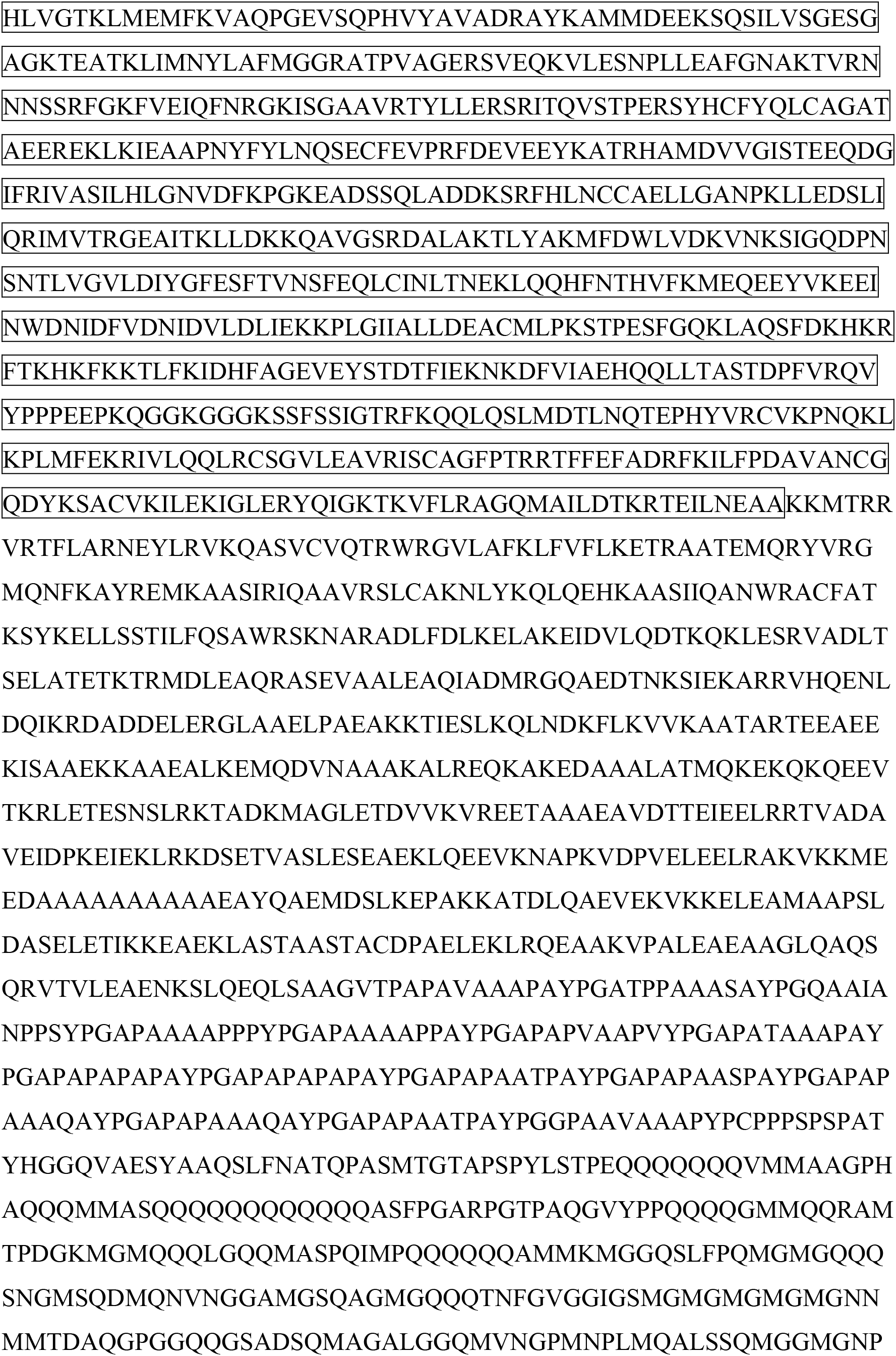
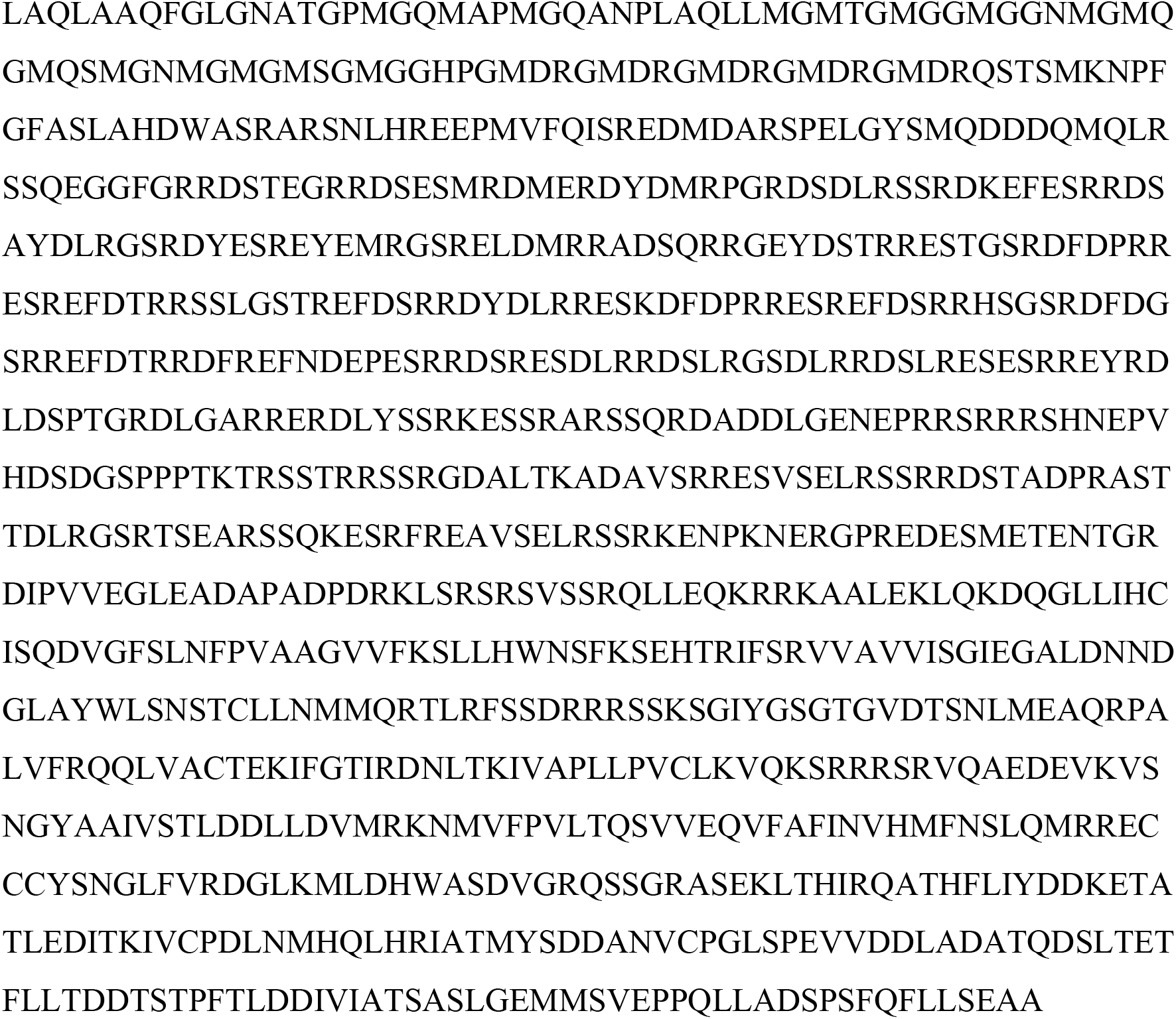
Amino acids sequence of *Cb*XI-1.

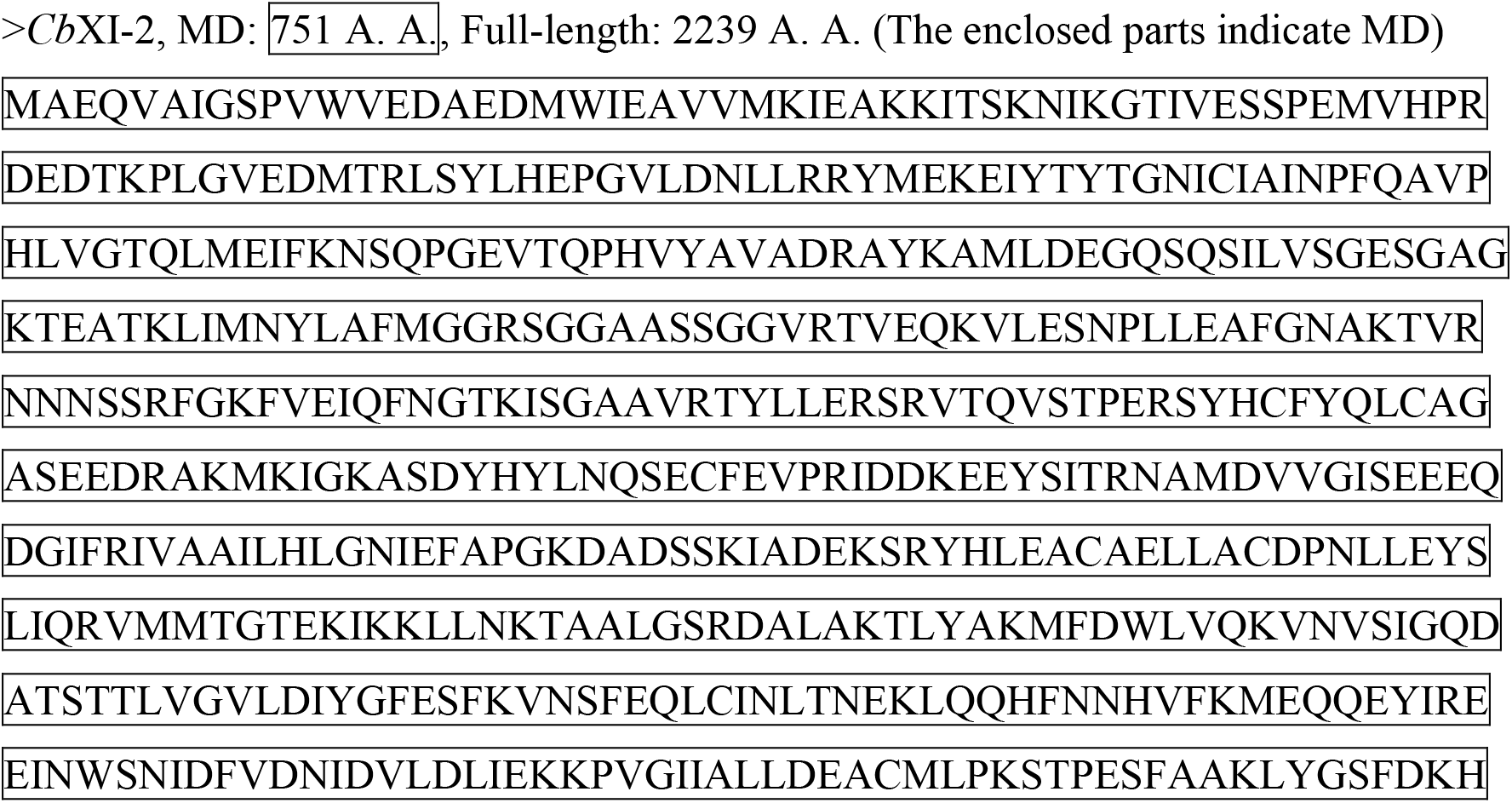
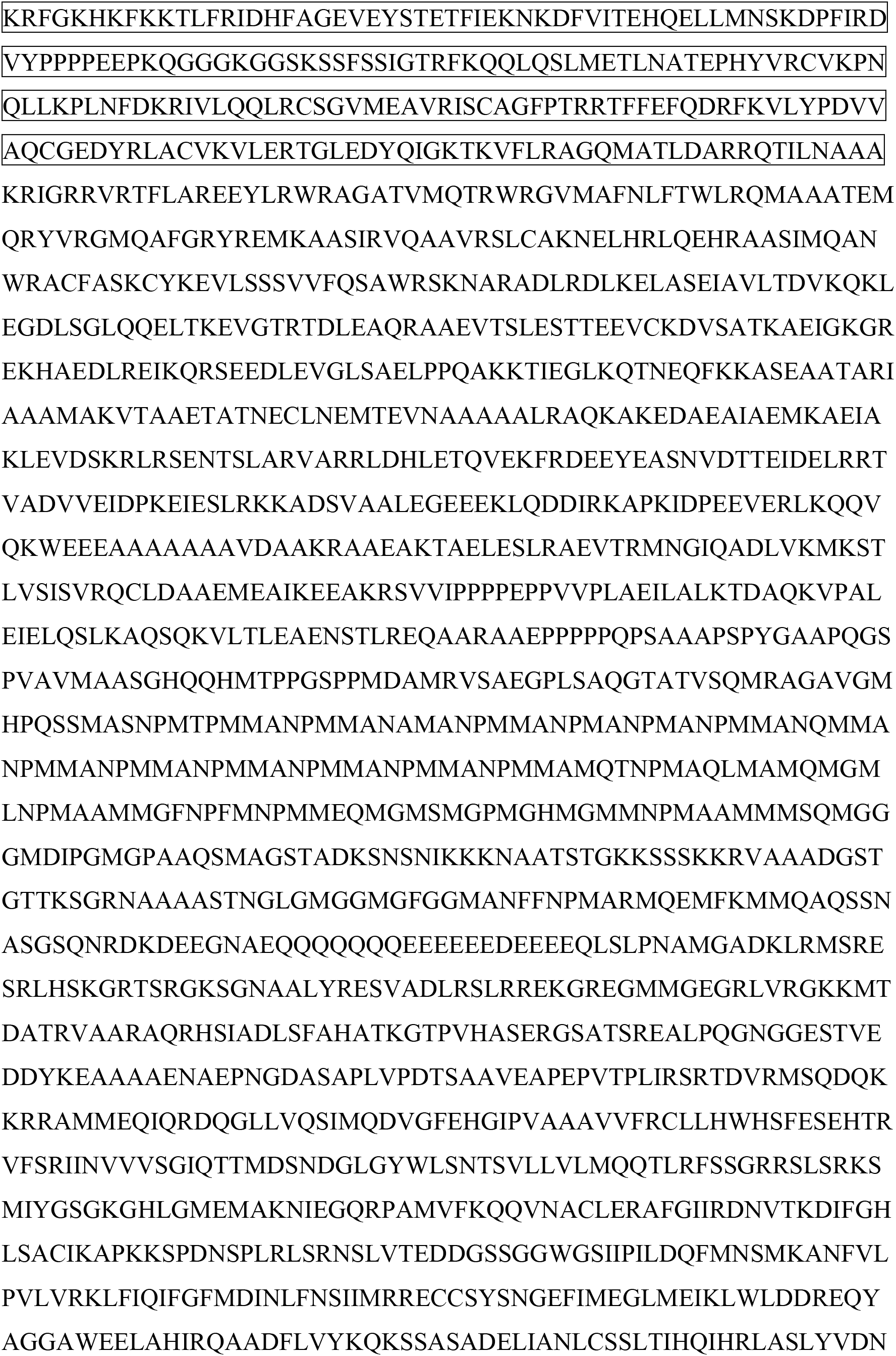
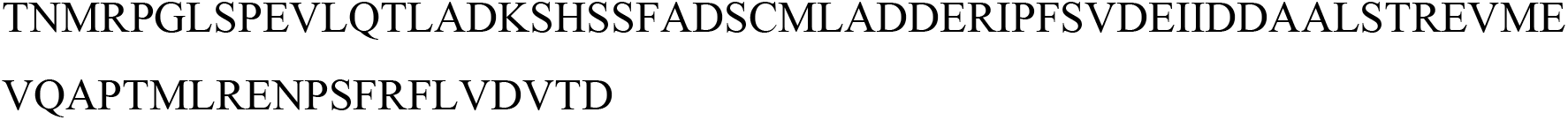
Amino acids sequence of *Cb*XI-2.

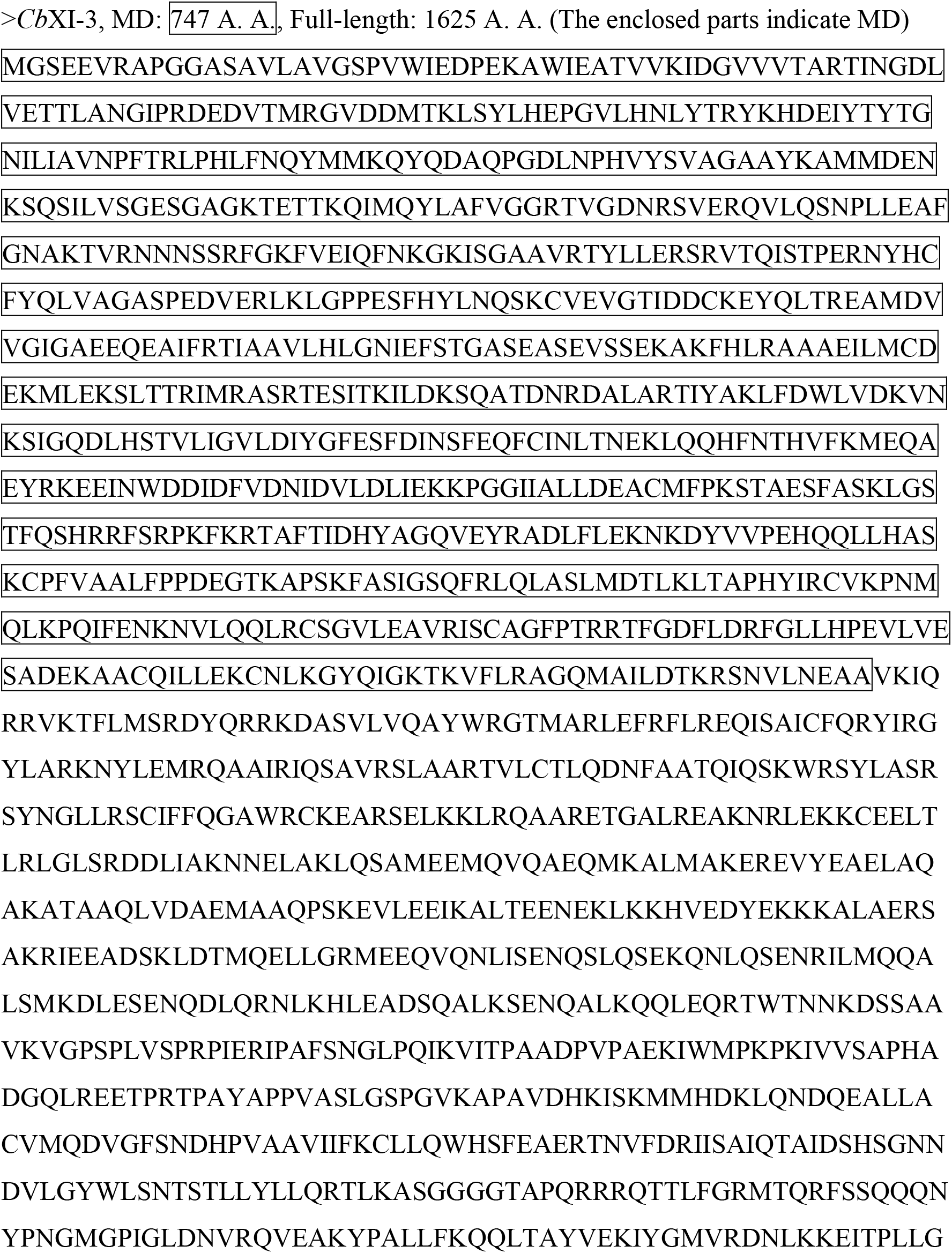
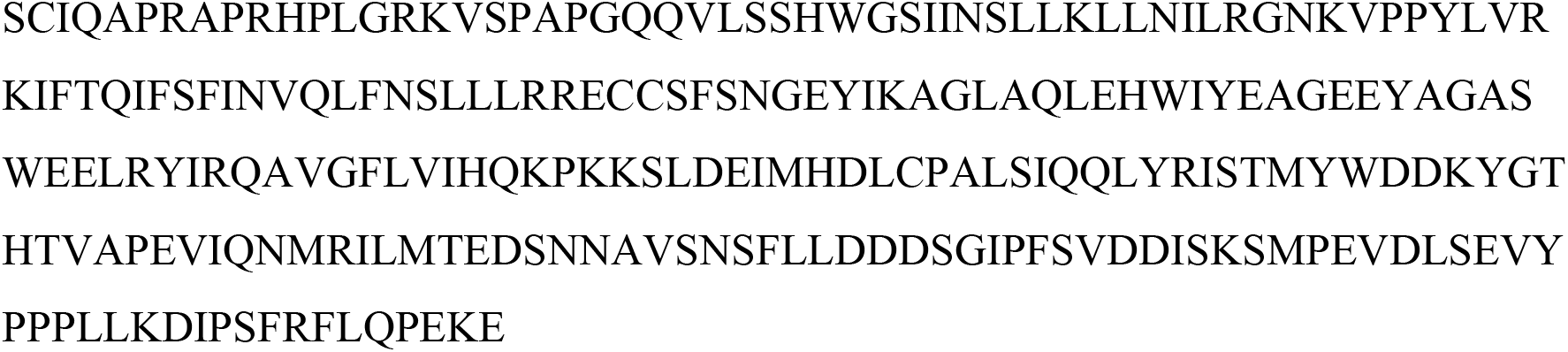
Amino acids sequence of *Cb*XI-3.

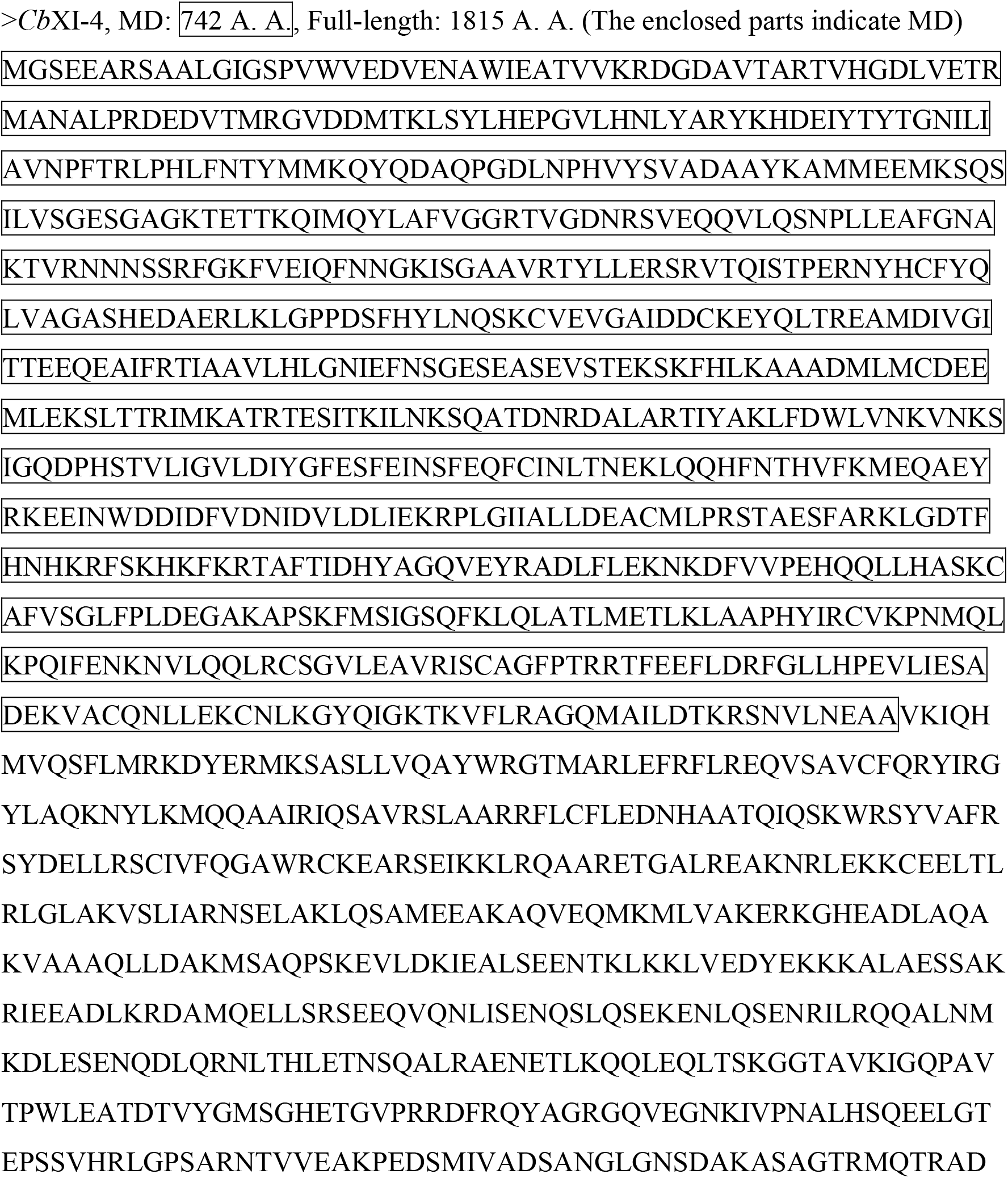
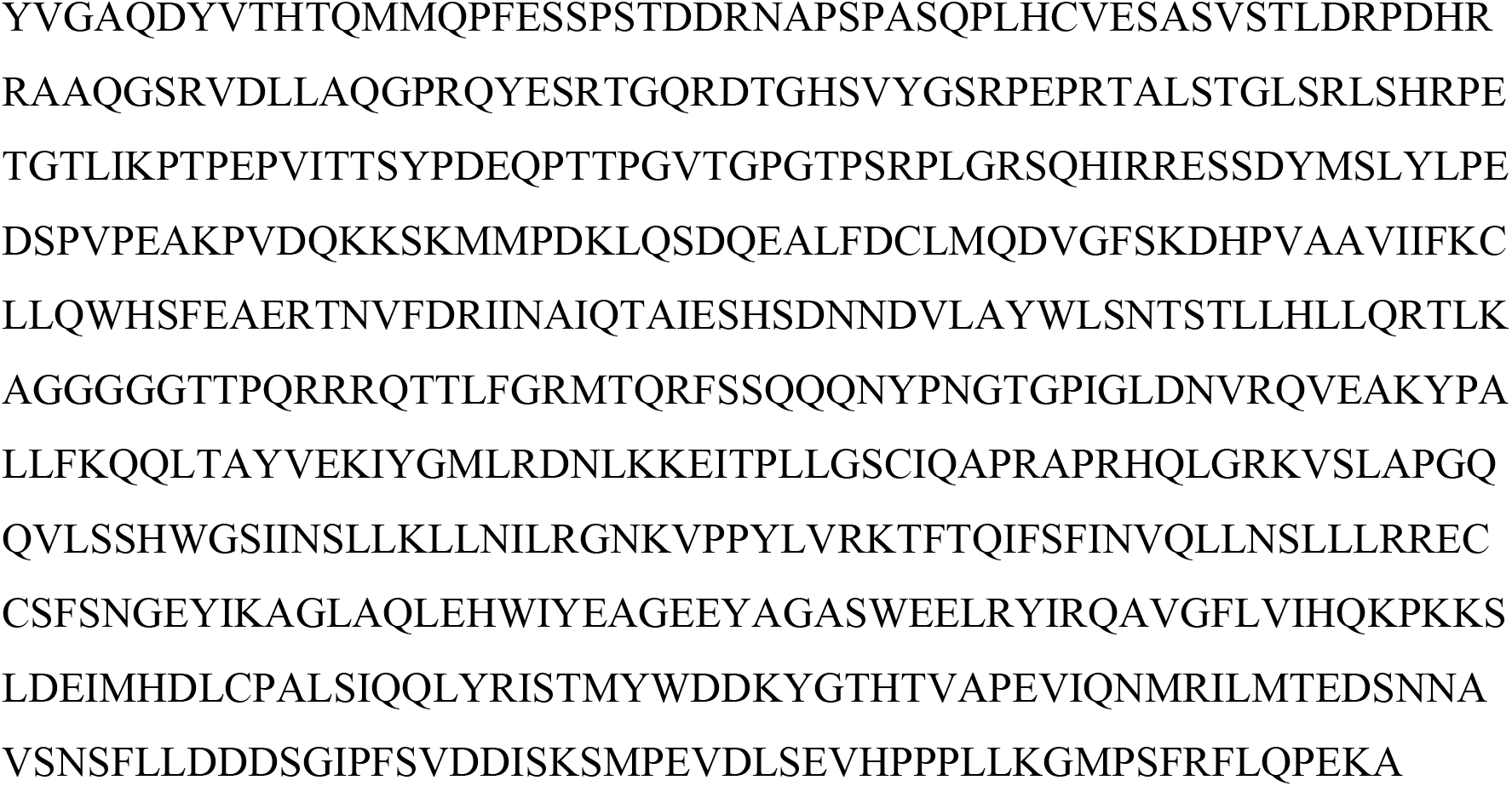
Amino acids sequence of *Cb*XI-4.

**Figure S1.**
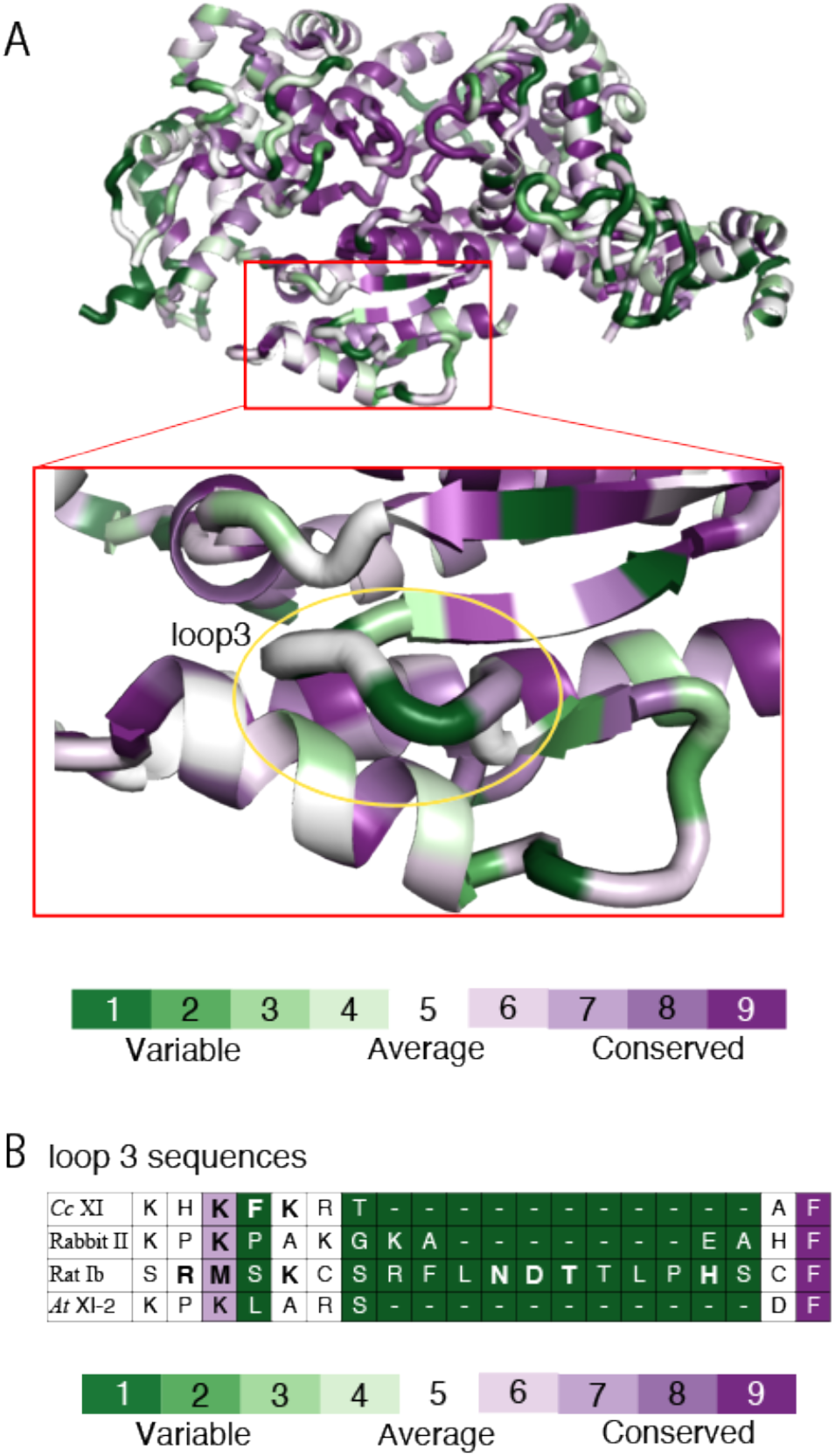
Actin-binding loop 3 with high diversity among myosins. (*A*) Amino acids of cryo-EM structure of *Cc* XI (23) are colored by conservation score ranging from green (1; most variable) to purple (9; most conserved residues), as shown in the color legend. The comparison of amino acid sequences was performed using *Cc* XI (23)(PDBID: 7KCH), Rabbit myosin II (24)(PDBID: 5H53), Rat myosin Ib (25)(PDBID: 6C1H) for which the cryo-EM structure of the actin-myosin complex have been reported, and *At* XI-2, which was successfully analyzed for crystal structure in this work. The viewing position, colors, and representations of the binding sites correspond to those in Fig.5B. (*B*) Alignment of amino acid sequence of loop 3 from *Cc* XI, Rabbit myosin II, Rat myosin IB, and *At* XI-2. In the cryo-EM structure of the actin-myosin rigor complex,amino acid residues with a distance of 4Å or less from actin are shown in bold.

**Table S1.**
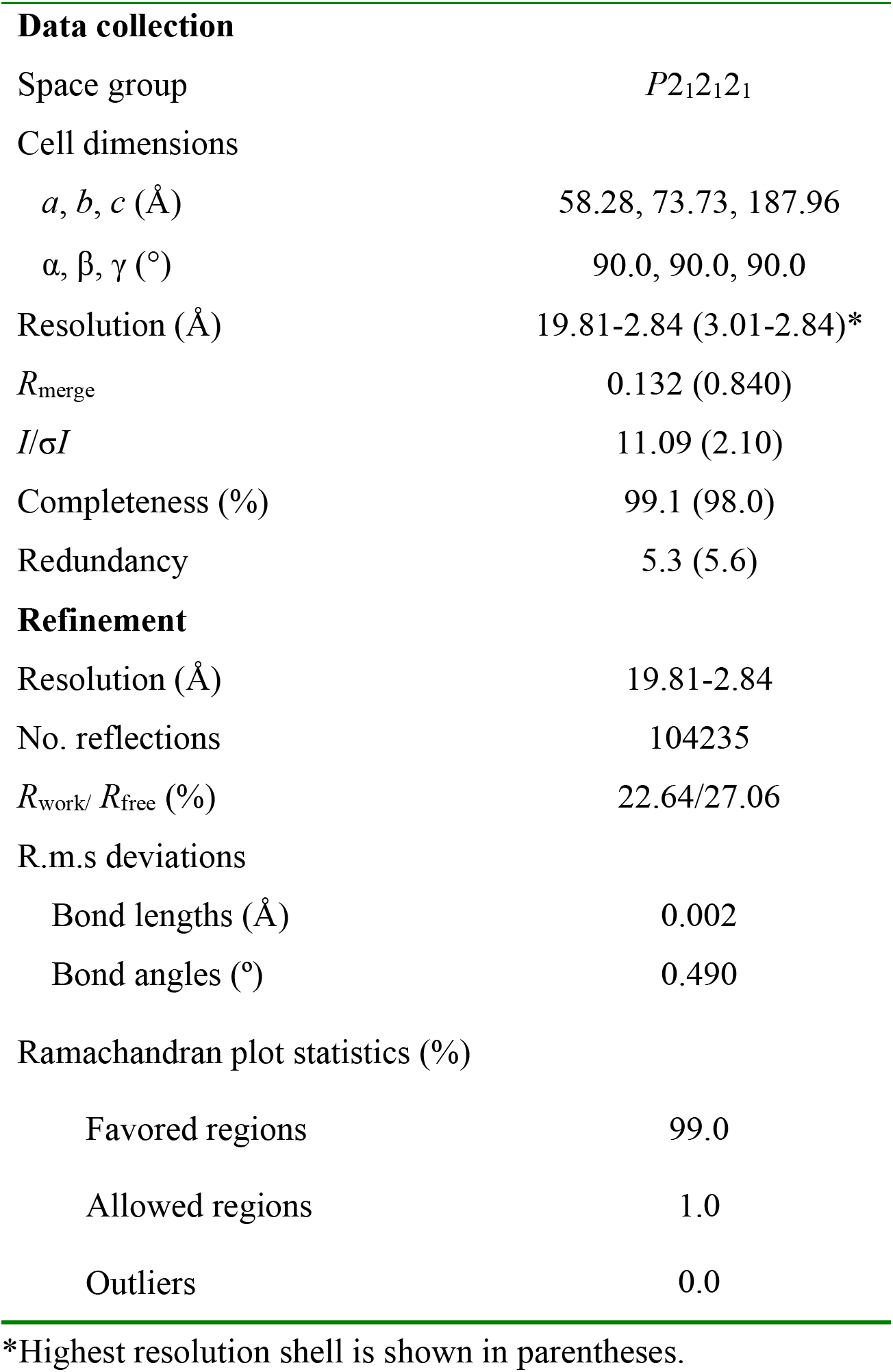
Crystallographic data collection and refinement statistics for *At* XI-2 MD

|  |  |
| --- | --- |
| **Data collection** |  |
| Space group | *P2<sub>1</sub>2<sub>1</sub>2<sub>1</sub>* |
| Cell dimensions |  |
| *a*, *b*, *c* (Å) | 58.28, 73.73, 187.96 |
| $\alpha$, $\beta$, $\gamma$ (°) | 90.0, 90.0, 90.0 |
| Resolution (Å) | 19.81-2.84 (3.01-2.84)* |
| *R*<sub>merge</sub> | 0.132 (0.840) |
| *I*/σ*I* | 11.09 (2.10) |
| Completeness (%) | 99.1 (98.0) |
| Redundancy | 5.3 (5.6) |
| **Refinement** |  |
| Resolution (Å) | 19.81-2.84 |
| No. reflections | 104235 |
| *R*<sub>work</sub>/ *R*<sub>free</sub> (%) | 22.64/27.06 |
| R.m.s deviations |  |
| Bond lengths (Å) | 0.002 |
| Bond angles (°) | 0.490 |
| Ramachandran plot statistics (%) |  |
| Favored regions | 99.0 |
| Allowed regions | 1.0 |
| Outliers | 0.0 |

\*Highest resolution shell is shown in parentheses.

**Table S2.** Expression levels of *Cb*XIs in *Chara braunii*

|  | RNA-Seq (FPKM) |  |  |  |
| --- | --- | --- | --- | --- |
|  | Whole plant | oogonia | antheridia | zygote |
| *CbXI-1* | 28.57 | 22.96 | 14.62 | 0.70 |
| *CbXI-2* | 3.96 | 15.17 | 6.13 | 13.59 |
| *CbXI-3* | 2.77 | 12.10 | 5.18 | 13.70 |
| *CbXI-4* | 3.95 | 8.42 | 8.33 | 2.94 |

data from (1)

## References

1. M. Kollmar, S. Muhlhausen, Myosin repertoire expansion coincides with eukaryotic diversification in the Mesoproterozoic era. BMC Evol Biol 17, 211 (2017).

2. M. A. Hartman, J. A. Spudich, The myosin superfamily at a glance. J Cell Sci 125, 1627–1632 (2012).

3. J. Walklate, Z. Ujfalusi, M. A. Geeves, Myosin isoforms and the mechanochemical cross-bridge cycle. J Exp Biol 219, 168–174 (2016).

4. T. Haraguchi et al., Molecular Characterization and Subcellular Localization of Arabidopsis Class VIII Myosin, ATM1. J Biol Chem 289, 12343–12355 (2014).

5. L. Golomb, M. Abu-Abied, E. Belausov, E. Sadot, Different subcellular localizations and functions of Arabidopsis myosin VIII. BMC Plant Biol 8, 3 (2008).

6. D. Van Damme, F. Y. Bouget, K. Van Poucke, D. Inze, D. Geelen, Molecular dissection of plant cytokinesis and phragmoplast structure: a survey of GFP tagged proteins. Plant J 40, 386–398 (2004).

7. A. Sattarzadeh, R. Franzen, E. Schmelzer, The Arabidopsis class VIII myosin ATM2 is involved in endocytosis. Cell Motil Cytoskeleton (2008).

8. T. Shimmen, E. Yokota, Cytoplasmic streaming in plants. Curr Opin Cell Biol 16, 68–72 (2004).

9. D. Avisar, M. Abu-Abied, E. Belausov, E. Sadot, Myosin XIK is a major player in cytoplasm dynamics and is regulated by two amino acids in its tail. J Exp Bot 63, 241–249 (2012).

10. M. Tominaga et al., Cytoplasmic streaming velocity as a plant size determinant. Dev Cell 27, 345–352 (2013).

11. V. V. Peremyslov, R. A. Cole, J. E. Fowler, V. V. Dolja, Myosin-Powered Membrane Compartment Drives Cytoplasmic Streaming, Cell Expansion and Plant Development. PLoS One 10, e0139331 (2015).

12. J. Verchot-Lubicz, R. E. Goldstein, Cytoplasmic streaming enables the distribution of molecules and vesicles in large plant cells. Protoplasma 240, 99–107 (2010).

13. K. Yamamoto et al., Chara myosin and the energy of cytoplasmic streaming. Plant and Cell Physiology 47, 1427–1431 (2006).

14. N. Kamiya, K. Kuroda, Velocity distribution of the protoplasmic streaming in Nitella cells. Bot Mag Tokyo 69, 544–554 (1956).

15. N. Kamiya, Protoplasmic streaming. In Handbuch der Planzenphysiologie (Ruhland, F., ed.), Springer, Berlin XVII, 979–1035 (1962).

16. R. E. Williamson, Actin in the alga, Chara corallina. Nature 248, 801–802 (1974).

17. K. Yamamoto, S. Hamada, T. Kashiyama, Myosins from plants. CMLS Cell. Mol. Life Sci. 56 (1999).

18. T. Q. Uyeda, Ultra-fast chra myosin: A test case for the swinging lever arm model for force production by myosin. J. Plant Res. 109, 231–239 (1996).

19. T. Kashiyama, N. Kimura, T. Mimura, K. Yamamoto, Cloning and characterization of a myosin from characean alga, the fastest motor protein in the world. J. Biochem. 127, 1065–1070 (2000).

20. M. Morimatsu et al., The molecular structure of the fastest myosin from green algae, Chara. Biochemical and Biophysical Research Communications 270, 147–152 (2000).

21. K. Ito et al., Recombinant motor domain constructs of Chara corallina myosin display fast motility and high ATPase activity. Biochem Biophys Res Commun 312, 958 – 964 (2003).

22. K. Ito et al., Kinetic mechanism of the fastest motor protein, Chara myosin. J Biol Chem 282, 19534–19545 (2007).

23. K. Ito, Y. Yamaguchi, K. Yanase, Y. Ichikawa, K. Yamamoto, Unique charge distribution in surface loops confers high velocity on the fast motor protein Chara myosin. Proc Natl Acad Sci U S A 106, 21585–21590 (2009).

24. T. D. Schindler, L. Chen, P. Lebel, M. Nakamura, Z. Bryant, Engineering myosins for long-range transport on actin filaments. Nat Nanotechnol 9, 33–38 (2014).

25. P. V. Ruijgrok et al., Optical control of fast and processive engineered myosins in vitro and in living cells. Nat Chem Biol https://10.1038/s41589-021-00740-7 (2021).

26. T. Nishiyama et al., The Chara Genome: Secondary Complexity and Implications for Plant Terrestrialization. Cell 174, 448–464 e424 (2018).

27. S. Kato et al., Morphology and Molecular Phylogeny of Chara altaica (Charales, Charophyceae), a Monoecious Species of the Section Desvauxia. CYTOLOGIA 75, 211–220 (2010).

28. S. Kato et al., New distributional records, taxonomy, morphology, and genetic variations of the endangered brackish-water species Lamprothamnium succinctum (Charales: Charophyceae) in Japan. Journal of Asia-Pacific Biodiversity 14, 15–22 (2021).

29. M. T. Casanova, K. G. Karol, A revision of Chara sect. Protochara, comb. et stat. nov (Characeae: Charophyceae). Australian Systematic Botany 27, 23–27, 25 (2014).

30. K. Yamamoto, M. Kikuyama, N. Sutoh-Yamamoto, E. Kamitsubo, Purification of actin based motor protein from *Chara corallina*. Proc. Japan Acad. 70, 175–180 (1994).

31. T. Q. P. Uyeda, P. D. Abramson, J. A. Spudich, The neck region of the myosin motor domain acts as a lever arm to generate movement. Proc. Natl. Acad. Sci. USA. 93, 4459–4464 (1996).

32. K. C. Holmes, The swinging lever-arm hypothesis of muscle contraction. Curr Biol 7, R112–118 (1997).

33. T. Haraguchi et al., Functional Diversity of Class XI Myosins in Arabidopsis thaliana. Plant Cell Physiol 59, 2268–2277 (2018).

34. M. Levitt, Conformational preferences of amino acids in globular proteins. Biochemistry 17, 4277–4285 (1978).

35. K. Imai, S. Mitaku, Mechanisms of secondary structure breakers in soluble proteins. Biophysics (Nagoya-shi) 1, 55–65 (2005).

36. M. Tominaga et al., Higher plant myosin XI moves processively on actin with 35 nm steps at high velocity. EMBO J 22, 1263–1272 (2003).

37. S. Muhlhausen, M. Kollmar, Whole genome duplication events in plant evolution reconstructed and predicted using myosin motor proteins. BMC Evol Biol 13, 202 (2013).

38. M. Kollmar, U. Durrwang, W. Kliche, D. J. Manstein, F. J. Kull, Crystal structure of the motor domain of a class-I myosin. EMBO J. 21, 2517–2525 (2002).

39. J. von der Ecken, S. M. Heissler, S. Pathan-Chhatbar, D. J. Manstein, S. Raunser, Cryo-EM structure of a human cytoplasmic actomyosin complex at near-atomic resolution. Nature 534, 724–728 (2016).

40. T. Fujii, K. Namba, Structure of actomyosin rigour complex at 5.2 A resolution and insights into the ATPase cycle mechanism. Nat Commun 8, 13969 (2017).

41. A. Mentes et al., High-resolution cryo-EM structures of actin-bound myosin states reveal the mechanism of myosin force sensing. Proc Natl Acad Sci U S A 115, 1292–1297 (2018).

42. Z. Duan, K. Ito, M. Tominaga, Heterologous transformation of Camelina sativa with high-speed chimeric myosin XI-2 promotes plant growth and leads to increased seed yield. Plant Biotechnology 37, 253–259 (2020).

43. A. Lupas, M. Van Dyke, J. Stock, Predicting coiled coils from protein sequences. Science 252, 1162–1164 (1991).

44. K. Collins, J. R. Sellers, P. Matsudaira, Calmodulin dissociation regulates brush border myosin I (110-kD-calmodulin) mechanochemical activity in vitro. J Cell Biol 110, 1137–1147 (1990).

45. T. Lin, N. Tang, E. M. Ostap, Biochemical and motile properties of Myo1b splice isoforms. J Biol Chem 280, 41562–41567 (2005).

46. T. Q. Uyeda, S. J. Kron, J. A. Spudich, Myosin step size. Estimation from slow sliding movement of actin over low densities of heavy meromyosin. J.Mol. Chem. 214, 699–710 (1990).

47. K. Ito, T. Q. Uyeda, Y. Suzuki, K. Sutoh, K. Yamamoto, Requirement of domain-domain interaction for conformational change and functional ATP hydrolysis in myosin. J Biol Chem 278, 31049 – 31057 (2003).

48. H. L. Sweeney et al., Kinetic tuning of myosin via a flexible loop adjacent to the nucleotide binding pocket. J. Biol. Chem. 273, 6262–6270 (1998).

49. F. Wang, E. V. Harvey, M. A. Conti, D. Wei, J. R. Sellers, A conserved negatively charged amino acid modulates function in human nonmuscle myosin IIA. Biochemistry 39, 5555 – 5560 (2000).

50. E. Golomb et al., Identification and characterization of nonmuscle myosin II-C, a new member of the myosin II family. J Biol Chem 279, 2800 – 2808 (2004).

51. S. Komaba, A. Inoue, S. Maruta, H. Hosoya, M. Ikebe, Determination of human myosin III as a motor protein having a protein kinase activity. J Biol Chem 278, 21352 – 21360 (2003).

52. T. Sakamoto, I. Amitani, E. Yokota, T. Ando, Direct observation of processive movement by individual myosin V molecules. Biochemical and Biophysical Research Communications 272, 586–590 (2000).

53. S. Watanabe, K. Mabuchi, R. Ikebe, M. Ikebe, Mechanoenzymatic characterization of human myosin Vb. Biochemistry 45, 2729 – 2738 (2006).

54. R. S. Rock et al., Myosin VI is a processive motor with a large step size. Proc. Natl. Acad. Sci. USA 98, 13655–13659 (2001).

55. I. P. Udovichenko, D. Gibbs, D. S. Williams, Actin-based motor properties of native myosin VIIa. J Cell Sci 115, 445–450 (2002).

56. P. L. Post, G. M. Bokoch, M. S. Mooseker, Human myosin-IXb is a mechanochemically active motor and a GAP for rho. J. Cell Sci. 111, 941–950 (1998).

57. K. Homma, J. Saito, R. Ikebe, M. Ikebe, Motor function and regulation of myosin X. J. Biol. Chem. 276, 34348–34354 (2001).

58. A. Herm_Gotz et al., Toxoplasma gondii myosin A and its light chain: a fast, single-headed, plus-end-directed motor. The Embo Journal 21, 2149–2158 (2002).

## References for Supplementary Information

1. T. Nishiyama et al., The Chara Genome: Secondary Complexity and Implications for Plant Terrestrialization. Cell 174, 448–464 e424 (2018).

2. K. Ito et al., Kinetic mechanism of the fastest motor protein, Chara myosin. J Biol Chem 282, 19534–19545 (2007).

3. M. Tominaga et al., Cytoplasmic streaming velocity as a plant size determinant. Dev Cell 27, 345–352 (2013).

4. K. Ito, Y. Yamaguchi, K. Yanase, Y. Ichikawa, K. Yamamoto, Unique charge distribution in surface loops confers high velocity on the fast motor protein Chara myosin. Proc Natl Acad Sci U S A 106, 21585–21590 (2009).

5. K. Ito et al., Recombinant motor domain constructs of Chara corallina myosin display fast motility and high ATPase activity. Biochem Biophys Res Commun 312, 958 – 964 (2003).

6. W. Kabsch, Xds. Acta Crystallogr D Biol Crystallogr 66, 125–132 (2010).

7. A. J. McCoy et al., Phaser crystallographic software. J Appl Crystallogr 40, 658–674 (2007).

8. P. Emsley, K. Cowtan, Coot: model-building tools for molecular graphics. Acta Crystallogr D Biol Crystallogr 60, 2126–2132 (2004).

9. P. D. Adams et al., PHENIX: a comprehensive Python-based system for macromolecular structure solution. Acta Crystallogr D Biol Crystallogr 66, 213–221 (2010).

10. S. C. Lovell et al., Structure validation by Calpha geometry: phi,psi and Cbeta deviation. Proteins 50, 437–450 (2003).

11. F.-W. Li et al., Fern genomes elucidate land plant evolution and cyanobacterial symbioses. Nature Plants 4, 460–472 (2018).

12. J. L. Bowman et al., Insights into Land Plant Evolution Garnered from the Marchantia polymorpha Genome. Cell 171, 287–304.e215 (2017).

13. F.-W. Li et al., Anthoceros genomes illuminate the origin of land plants and the unique biology of hornworts. Nature Plants 6, 259–272 (2020).

14. S. Cheng et al., Genomes of Subaerial Zygnematophyceae Provide Insights into Land Plant Evolution. Cell 179, 1057–1067.e1014 (2019).

15. K. Katoh, D. M. Standley, MAFFT multiple sequence alignment software version 7: improvements in performance and usability. Mol Biol Evol 30, 772–780 (2013).

16. W. P. Maddison, D. R. Maddison, Mesquite: a modular system for evolutionary analysis. Version 3.61 http://www.mesquiteproject.org(2019).

17. A. Stamatakis, RAxML version 8: a tool for phylogenetic analysis and post analysis of large phylogenies. Bioinformatics 30, 1312–1313 (2014).

18. J. Felsenstein, PHYLIP (Phylogeny Inference Package) version 3.697. (Distributed by the author. Department of Genome Sciences, University of Washington, Seattle.). (2017).

19. S. F. Altschul et al., Gapped BLAST and PSI-BLAST: a new generation of protein database search programs. Nucleic Acids Res 25, 3389–3402 (1997).

20. G. Robertson et al., De novo assembly and analysis of RNA-seq data. Nature Methods 7, 909–912 (2010).

21. M. Stanke et al., AUGUSTUS: ab initio prediction of alternative transcripts. Nucleic Acids Research 34, W435–W439 (2006).

22. E. Lee et al., Web Apollo: a web-based genomic annotation editing platform. Genome Biology 14, R93 (2013).

23. P. V. Ruijgrok et al., Optical control of fast and processive engineered myosins in vitro and in living cells. Nat Chem Biol 10.1038/s41589-021-00740-7 (2021).

24. T. Fujii, K. Namba, Structure of actomyosin rigour complex at 5.2 A resolution and insights into the ATPase cycle mechanism. Nat Commun 8, 13969 (2017).

25. A. Mentes et al., High-resolution cryo-EM structures of actin-bound myosin states reveal the mechanism of myosin force sensing. Proc Natl Acad Sci U S A 115, 1292–1297 (2018).

